# Heat Stress on the brown Seaweed *Ascophyllum nodosum*: differential population sensitivity to future climate

**DOI:** 10.1101/2024.10.14.618208

**Authors:** Luís F. Pereira, Francisco Arenas, Rui Seabra, Rita da Silva, Cátia Monteiro, Joana Pereira, Cândida G. Vale, João Serôdio, Silja Frankenbach, Vittoria Ghiglione, Pedro Ribeiro, Fernando P. Lima

**Affiliations:** CIBIO, Centro de Investigação em Biodiversidade e Recursos Genéticos, InBIO Laboratório Associado, Campus de Vairão, Universidade do Porto, 4485-661 Vairão, Portugal; BIOPOLIS Program in Genomics, Biodiversity and Land Planning, CIBIO, Campus de Vairão, 4485-661 Vairão, Portugal; Departamento de Biologia, Faculdade de Ciências, Universidade do Porto, 4099-002 Porto, Portugal; Benthic Ecology and Environmental Solutions Team, CIIMAR, University of Porto, Terminal de Cruzeiros do Porto de Leixões, Av. General Norton de Matos s/n, 4450-208 Matosinhos, Portugal; CESAM — Centre for Environmental and Marine Studies and Department of Biology, University of Aveiro, 3810-193 Aveiro, Portugal; Department of Biological Sciences, University of Bergen, 5006 Bergen, Norway

**Keywords:** *Ascophyllum nodosum*, Habitat-forming species, Rocky shore, Intertidal thermal stress, Population sensitivity, Climate change

## Abstract

Accurate forecasts of the biological impacts of climate change require a better understanding of how small-scale temperature variability affects individual physiology and population dynamics. But doing so for intertidal species with large distribution ranges, while also accounting for the effect of local adaptation, presents numerous technical challenges. Here, we assessed the macroecological consequences of thermal stress on the cold-adapted brown seaweed *Ascophyllum nodosum* across its European distribution. We collected specimens from ten populations spanning latitudes 41°N to 60°N and subjected them to simulated intertidal heat stress using a novel, custom-built experimental setup that replicated realistic conditions, including tidal cycles, light conditions, and temperature trajectories based on *in situ* data. Our factorial design comprised eight experimental treatments, combining two high-tide water temperatures (15 °C and 20.5 °C) with four low-tide peak temperatures (28.5 °C to 40.5 °C). Physiological performance was evaluated through measurements of growth, mortality, and oxygen production. Results indicate that thermal stress is more closely associated with the magnitude of temperature change between high and low tides rather than the absolute maximum temperatures reached. Algae exposed to warmer water temperatures (20.5 °C) consistently outperformed those in colder water (15 °C), suggesting that cold upwelled waters at the species’ southern limit may not be essential for survival. Southern populations demonstrated higher resilience to thermal stress than central and northern counterparts. Integrating these physiological responses with climate projections, we employed demographic models to forecast long-term population dynamics. The models predict that future climatic conditions could exceed the thermal resilience of specific populations, leading to uneven impacts across the European distribution of the species. Notwithstanding, range contractions may occur at the warm edge of the distribution, where populations, though more resilient to thermal stress, could still be overwhelmed by the pace of warming. This study underscores the importance of realistic experimental simulations in evaluating species’ thermal tolerance and highlights the potential for climate change to differentially impact populations along large latitudinal gradients.

## Introduction

Temperatures have been following alarming trends in both atmospheric and marine domains (Kemp et al., 2022; Oliver et al., 2019). These trends are expected to intensify, with global air temperatures predicted to rise between 1 °C and 5 °C and sea surface temperatures between 1.6 °C and 4 °C until the end of the century (Pörtner et al., 2022). Importantly, since warming patterns differ among the two domains, temporal and spatial heterogeneity at smaller scales may also be on the rise (Coumou et al., 2013). Understanding how this small-scale temperature variability affects the individual’s physiology and subsequent population resilience is, therefore, critical for accurately predicting the biotic impacts of climate change (Choi et al., 2019). Species inhabiting intertidal rocky shores provide exceptional opportunities for these studies. Despite their marine origin, they periodically endure terrestrial conditions during low tide, when body temperatures can fluctuate by as much as 20 °C in a couple of hours (Seabra et al., 2011). Consequently, rocky shore organisms are considered sensitive indicators of climate change (Mislan et al., 2014; Wethey & Woodin, 2008). Yet, the explicit incorporation of the thermal complexity of intertidal zones within experimental studies presents substantial challenges.

First, the identification of appropriate experimental temperature treatments isn’t always trivial. A common framework is identifying heatwaves and cold snaps, defined as significant deviations from typical climate conditions (Domeisen et al., 2023; Hobday et al., 2016), and using them to characterize and quantify weather anomalies. But although extreme events are evidently correlated with mass mortality events among algae (Thomsen et al., 2019), seagrass (Strydom et al., 2020), and benthic invertebrates (Le Nohaic et al., 2017; Traiger et al., 2022), mild but realistic temperature changes can also lead to meaningful impacts. For example, survivability in mussels (*Perna canaliculus*) following field transplants was found to be correlated with time spent in not-so-extreme heat (>25°C) during emersion periods (Benjamin et al., 2024), a detail that could only be properly captured with *in situ* temperature sensors. Additionally, these impacts can be more severe when habitat-forming species are affected, triggering cascading effects on biodiversity, community structure, and ecosystem services (Garrabou et al., 2022; Hesketh & Harley, 2023; Smale, 2017).

Second, environmental data must be acquired at scales relevant to the organisms under study (Potter et al., 2013), and that can be very hard to accomplish. Obtaining long-term and accurate data for the coastal fringe is already difficult during high tides (Brewin et al., 2018; Castillo & Lima, 2010) because remotely sensed data often lack the spatial and temporal resolution needed to capture sharp gradients in sea temperature near the coast (Meneghesso et al., 2020). This task becomes even more challenging during low tides when even data from nearby weather stations have been shown to overlook extreme variations in intertidal air temperature (Lathlean et al., 2011). However, ignoring those temperatures altogether may result in the mischaracterization of critical thermal limits for some species. Crucially, while still a logistically complex task, this challenge has now been largely resolved through the development of small, sturdy, and autonomous temperature loggers (e.g., Lima and Wethey (2009)) that can consistently record hourly temperature data over extended periods of time and at the scale of the study organisms both during high and low tides (Lima et al., 2016).

Third, the potential geographical variability in the physiological responses of the organisms themselves must be accounted for. This is particularly concerning in species with broad geographic ranges and low mobility (Straub et al., 2019), such as macroalgae, which are likely to exhibit phenotypic plasticity or local adaptation (Hernández et al., 2023; King et al., 2018). One of the most usual methods to detect population-level variability in heat tolerance involves conducting common-garden experiments where populations from different origins are tested under uniform environmental conditions (King et al., 2018). So far, results have varied across species and experiments, with findings sometimes pointing to heat-adapted distribution edges and other times uncovering unexpectedly tolerant central populations (DuBois et al., 2022; King et al., 2019). Regardless, it is important to recognize that even heat-tolerant populations may face future challenges if the temperatures they experience eventually exceed their adaptive thresholds (Pearson et al., 2009). Thus, the significance of a population’s heat resilience can only be fully understood within the context of the projected climate it will endure.

With these challenges in mind, we aimed to assess the macroecological consequences of the physiological responses of the fucoid *Ascophyllum nodosum* (Fucales, Phaeophyeceae) to simulated intertidal heat stress across its distributional range. *A. nodosum* is an important habitat-forming species present along both North Atlantic coastlines, commonly found in sheltered intertidal rocky shores from Norway to Portugal in the East and from Baffin Island, Canada to New York, USA in the West. It is a relatively slow-growing, long-lived species with an estimated maximum lifespan of 60 years, and it is capable of producing clonal individuals through vegetative sprouting from its base (Åberg, 1992b). Commonly referred to as a cold-adapted species, summer temperatures are thought to be a limiting factor for its southern populations (Lüning, 1991). Importantly, these southern populations are not contiguous to the rest of the species range. Instead, they are geographically separated and restricted to the NW corner of the Iberian Peninsula, a biogeographic stronghold for cold-adapted species due to the upwelling of cold water during the summer months (Monteiro et al., 2022).

We used a custom-built experimental set-up to realistically simulate tides, light, and, importantly, daily submersion and emersion temperature trajectories through the fine control of water temperature, infrared (IR) radiation, and air circulation. Sampled populations span a latitudinal range from 41° to 60° N, encompassing most of the species’ latitudinal range across Europe. Realistic temperature trajectories were based on hourly temperature data from *in situ* loggers deployed in the sampled populations. To understand the importance of cold water in the upwelling-dominated southern distribution limit, we simulated two water temperature scenarios, one typical of a period of strong upwelling (colder) and another corresponding to upwelling relaxation (warmer). We then evaluated physiological performance through growth, mortality, and oxygen production. From the collected data, we inferred evidence of local adaptation to thermal stress. We combined current population-specific physiological responses with future temperature projections and ran demographic models (David R. Schiel, 2013) parameterized to the specific populations under study (Åberg, 1992b) so as to produce long-term population forecasts across the European range of this species.

## Materials and methods

### Sample collection

Juvenile vegetative fronds of *Ascophyllum nodosum* measuring 6 to 10 cm in length were collected between 15 and 25 of August 2021 at low tide from ten populations (Fig. 1a; Supplementary Table S1) across the species’ vertical distribution and capturing most of its European latitudinal range (Pereira et al., 2020). Sampling avoided nearby individuals, minimizing the chance of collecting clones. After collection, fronds were transported to the Interdisciplinary Centre of Marine and Environmental Research (CIIMAR, Porto, Portugal) within 24 hours in refrigerated boxes. Upon arrival, they were acclimated separately for two weeks at constant aeration and at 150-175 µmol photon m^-2^ s^-1^ PAR irradiance (US-SQS/L PAR sensor, WALZ, Germany) with 12h/12h photoperiod (light/dark) under 5032 LED stripes. Acclimation followed an intertidal cycle of 7/5 hours (high/low), simulating the observed tides at the vertical niche of *A. nodosum* in Viana do Castelo (Portugal). During acclimation, water temperature was slowly brought from the water temperature observed at the collection site to the targeted treatment at a daily maximum rate of 1 °C.

**Fig. 1.**
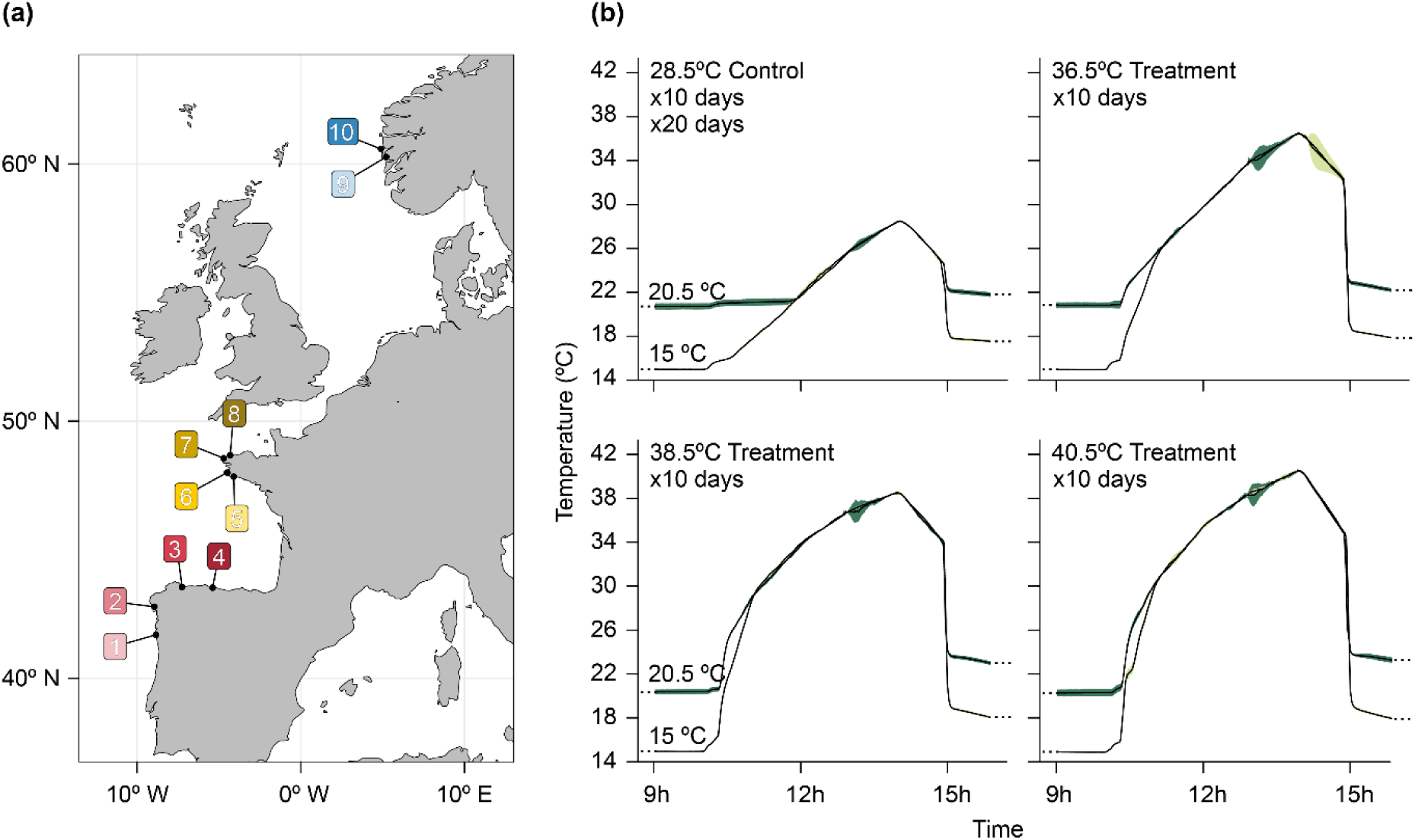
(a) Map of the study area. Numbers 1–10 indicate the locations of the ten populations sampled (1– Viana do Castelo, PT; 2-Ría de Muros, ES; 3-Ría de Foz, ES; 4-Ría de Villaviciosa, ES; 5-Île-Tudy, FR; 6-Penmarch, FR; 7-Landunvez, FR; 8-Soulougan, FR; 9-Blomsterdalen, NO; 10-Straumøyna, NO). (b) Average diurnal low tide temperatures within each experimental tank for the 8 treatments (black line) ± SD (light green and dark green for cold– and warm-water treatments, 15 °C and 20.5 °C, respectively). Nocturnal low tide temperatures remained close to their respective water temperatures. The tide cycle of 7:5 (high: low) was repeated twice daily for the whole 20 days of the experiment. Map created in R (R Development Core Team, 2023) using the Rnaturalearth library v1.0.0.

### Experimental design

The experiment (Fig. 1b) was designed to provide short-term exposure to heat stress during low tides while simulating realistic tidal cycles with two different high tide temperatures. A 7/5 h (high/low) tide cycle was applied to all treatments during the whole experiment. The experimental design had four low tide and two high tide temperatures, composing a full factorial design of 4×2 levels. Water temperature during the experiment was fixed at 15 °C (cold) or 20.5 °C (warm), depending on the treatment. These water temperatures correspond to the summer average values at the southern edge population (Viana do Castelo, Portugal) during strong and weak upwelling events, respectively, and were obtained from an hourly *in situ* temperature dataset collected between 2010 and 2020. Although seasonal upwelling keeps seawater around 15 °C during most of the summer, occasional relaxation of upwelling promotes warming periods, providing insight into the conditions of a weakened upwelling system.

Low tide periods were characterized by emersion accompanied by realistic heating simulated with infrared-emitting lamps. To assess the effect of an acute heatwave on *A. nodosum* performance, we selected the most extreme hot days from the temperature record (i.e., those days whose temperatures exceeded the overall 95^th^ percentile). We then determined that peak temperatures were typically reached four hours after the emersion began and calculated the average temperature trajectory slopes leading to and from this peak point. These data were used to generate four diurnal low-tide temperature profiles with similar shapes but reaching different temperature peaks (Fig. 1b). The lowest peak, set at 28.5 °C, was used as the control treatment and corresponded to the average summer temperature at the southern edge population. The other three experimental peak temperatures were chosen based on preliminary tests that indicated a peak temperature of approximately 38.5 °C as a potential tipping point for *A. nodosum’*s growth performance. To capture any critical tipping performance point, which could occur at slightly lower or higher temperatures due to the broader selection of populations under test, we added the peak temperature treatments of 36.5 °C and 40.5 °C alongside the 38.5 °C treatment. In summary, the experiment involved eight treatments in a 4×2 factorial design, combining four low-tide peak temperatures (28.5 °C, 36.5 °C, 38.5 °C, and 40.5 °C) with two high-tide water temperatures (15 °C and 20.5 °C). These eight temperature profiles (Fig. 1b) were applied for 10 consecutive days, followed by an additional 10-day period during which all experimental conditions were set to the control treatment, allowing any delayed stress or recovery effects to manifest.

### Experimental common-garden system

The common-garden system was composed of 16 polycarbonate 30-liter tanks (n = 2 per treatment), maintained in a water bath system to regulate water temperatures. Water bath temperature was controlled using a combination of three 500 W titanium heaters and a water chiller (TECO TR15). Room temperature was maintained at 17 °C to reduce temperature fluctuations during nocturnal low tides. Light was kept constant at 150-175 µmol photon m^-2^ s^-1^ PAR in a 12-hour light/dark photoperiod cycle. Tide control was achieved via automatization of valves (U.S. Solid, USA). At low tide, seawater was discarded from each tank, and at high tide, new seawater was pumped in. Separated filling system, complete water replacement during each tidal cycle, and continuous aeration ensured independent systems and prevented water deterioration effects. Low tide temperatures were controlled using two 125 W IR lamps per experimental tank, regulated by a customized Proportional-Integral-Derivative (PID) algorithm. To prevent temperature stratification during low tide, a 60 mm wide fan with 43.9 m³ h^-1^ air flow was turned at low tides. Two temperature loggers (Lima & Wethey, 2009) were deployed inside each tank attached to granite tiles, providing temperature feedback to a microcontroller (ATMEGA2560), which coordinated all electrical functions (Seabra et al., 2016). All algae were identified with wire markers and held on the granite tile surface with elastic fabric cord (10 individuals per population on each tank; n=20 treatment^-1^).

### Relative growth rates

Relative growth rates (RGR), expressed as the percentage change in fresh weight (FW) per day (Hurd et al., 2014) were calculated on the day 10 and day 20 of the experiment (n = 20 per population).

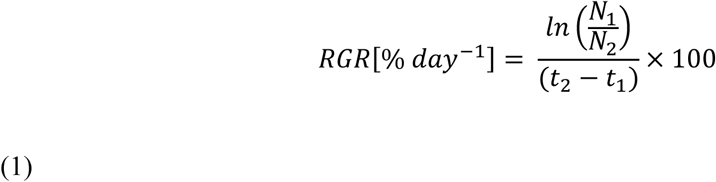

Where *N_1_* and *N_2_* represent the initial and final fresh weight, respectively, and *t_1_* and *t_2_* are the time points in days. Only fronds determined as whole and alive were considered in RGR results. Fragmented fronds, although they could be considered alive, were not reliable for growth measurements. For measurement, individuals were briefly dried on paper cloth before weighing with a 0.01 g precision scale (Kern EG 220-eNM, Mechanikus Gottlieb KERN, Germany).

### Survival

Survival was determined by chlorophyll fluorescence imaging on day 10 and day 20 of the experiment. Individuals were considered dead if their images showed no pixels with fluorescence above the background level. Chlorophyll fluorescence was measured for all individuals using a FluorCAM 800MF imaging fluorometer (Photon System Instruments, Brno, Czechia), as described by Serôdio and Campbell (2021). Images of chlorophyll fluorescence parameters (dark-level fluorescence) were captured for all seaweeds after a minimum of 20 minutes of dark adaptation, followed by a modulated measuring light and saturation pulses (< 0.1 and > 7500 μmol photons m-2 s-1, respectively), provided by red LED panels (612 nm emission peak, 40-nm bandwidth). Images (512×512 pixels) were analysed using FluorCam7 software (Photon System Instruments, spol. s r.o., Drasov, Czechia). Areas of interest (AOIs) were defined to match the whole area of each seaweed, excluding the non-fluorescent background signal.

### Primary productivity

Primary productivity was quantified in all treatments at day 10 and day 20 of the experiment by measuring oxygen production and respiration. Fronds (n = 6 per population) were placed in 60 ml acrylic chambers filled with filtered seawater (1 μm polypropylene filter) and kept in a water bath at the respective treatment temperature (15 °C or 20.5 °C) for productivity-irradiance curve analysis (P-I curves). Each chamber contained a magnetic stirring bead to ensure seawater homogenization, separated by an acrylic mesh to avoid damaging the algae. To measure oxygen production and consumption (respiration), light intensity followed a gradient of 0 (dark), 50, 180, 330, 550, and 1000 µE m^-2^ s^-1^. Light intensity was measured with a calibrated quantum irradiance meter (US-SQS/L, ULM-500; Waltz GmbH, Effeltrich, Germany). Light intensities were obtained from cool white LED strips (model 5032), assembled in five groups, and controlled by an ATMEGA 328p microcontroller with a 5-relay setup. Light periods started with 15 minutes of darkness, followed by sequential 10-minute steps at each light intensity. The water volume in each chamber, used for posterior oxygen rate calculations, was specific to each individual, with the frond’s volume subtracted from the total chamber volume. Oxygen content was measured once per minute by an optical fiber probe (DP-PSt7, Oxy-4 SMA, PreSens Precision Sensing GmbH, Germany). Productivity measurements were conducted in sets of eight chambers at a time, with one being a control blank chamber. Oxygen rates were calculated in µmol O^2^ ww g^-1^ h^-1^ in 5-minute intervals. Dark respiration (R_d_) and net primary productivity (NPP) rates were estimated as the slope of the linear regression between oxygen concentration and time. Gross photosynthesis (GP) was calculated by adding the absolute value of R_d_ to the NPP estimates at each light intensity. The maximum rate of gross photosynthesis at saturating light (GP_max_) was determined by fitting the model of Eilers and Peeters (1988):

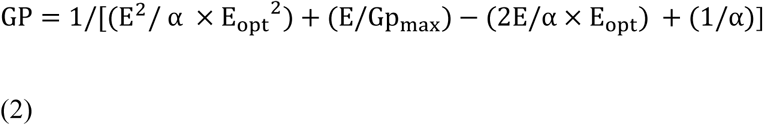

Where E is irradiance, E_opt_ is the optimum irradiance, and α is photosynthetic efficiency at low irradiance. The maximum rate of net primary production (NPP_max_) was calculated as GPmax – R_d_. Model fitting was performed by non-linear regression using the Phytotools package for R (Silsbe & Malkin, 2015).

### Statistical analysis

We used generalized linear mixed models (GLMMs; Supplementary Table S6) with a Gaussian distribution to analyse the effect of each treatment on growth and oxygen production (Zuur et al., 2009). Analyses were made separately for the results on day 10 and day 20. The best model for each parameter was found by gradual simplification of the full model until the lowest Akaike Information Criterion (AIC) was found (Burnham & Anderson, 2004). The full model included high-tide temperature (two levels: 15 °C and 20.5 °C), low-tide temperature (four levels: 28.5 °C, 36.5 °C, 38.5 °C, and 40.5 °C), and region (three levels: South, Center, and North) as fixed factors. The population was included as a random effect nested within the region, and the tank was included as a cross-random effect. Model normality was checked by visual inspection of Q-Q plots, and heteroscedasticity was assessed through residual plots. Random effects showed negligible contributions to the model’s variance for all the models. To investigate pairwise differences between treatment groups, we used estimated marginal means (emmeans). A significance threshold of 0.05 was used for testing treatment effects. The survival rate was analysed using GLMM as well but with binomial distribution with a logit-link function. For direct analysis of survival after 10 days, the factors low– and high-tide temperatures were replaced by a continuous variable representing the temperature difference between the two (8 °C to 26.5 °C). While population survival models used in forecasting mortalities followed the lower AIC method described for growth for the simplest model (Supplementary Table S2). Statistical analyses were done in R4.3.1 (R Core Team, 2023), with the lme4 (v1.1-35.1) and the emmeans (v1.8.7) (Lenth et al., 2021) packages.

### Modelling strategy

To predict survival and population size as a consequence of the physiological responses observed here, we required temperature data for low and high tides that would be analogous to *in situ* measurements. Although *in situ* measurements are accurate, their availability is limited in both space and time due to substantial costs and logistical challenges involved in data collection. Additionally, these measurements are influenced by local-scale factors, making it difficult to extrapolate them to future climate scenarios. To address this limitation, we developed a model relating *in situ* daily maximum temperature of intertidal rock (*T*_max_) to weather model data, and used it to obtain forecasts of low tide maximum temperatures from fine-scale climatic data.

### Forecasting *in situ* data

*In situ* data was obtained from autonomous temperature loggers (Envloggers, Electricblue,CRL, Portugal) deployed at the center of the tidal range of *A. nodosum* at ten population sites for 2 years (August 2021 to October 2023). Temperature data were collected hourly at 0.1 °C resolution (n = 2 per site). Loggers were deployed in 30-mm wide and 15-mm deep drilled holes and sealed with marine epoxy (Z-Spar™ Splash Zone Compound A-788). All sensors were facing south on ∼45° angled rocks for maximum sun exposure to effectively capture the low tide maximum rock temperature.

Weather data were gathered from ERA5 reanalysis datasets (Hersbach, 2023; Muñoz-Sabater et al., 2021), produced by the European Centre for Medium-Range Weather Forecasts (ECMWF). Variables included 2 m temperature (t2m), 10 m u and v wind components (10u and 10v, respectively), total precipitation (tp), and sea surface temperature (sst). All variables were provided as hourly values with spatial resolutions of 0.1 x 0.1 arc degrees (with the exception of sst, available as a 0.25 arc degree grid). Data pre-processing involved summarizing temperature to daily maximum for both *in situ* loggers and ERA5 t2m (t2m_max_), calculating the daily sst average (sst_mean_), converting total precipitation from m h^-1^ to mm s^-1^ (tp_mean_), and converting wind speed from scalar to magnitude by applying the square root of the sum of the squares of its u and v components (wind_mean_). All climatic variables were accessed through Copernicus Climate Data Store (CDS), at coordinates closest to each of the ten sites (Supplementary Table S3). Environmental variables for predicting temperature were selected based on their availability across the reanalysis and forecasted datasets (ERA5 re-analysis and CMIP6 projections) and their informative value for the model. Since violations of linear model assumptions can compromise reliable predictions, we used a Generalized Least Square (GLS) model, which provides more efficient estimates when the residuals exhibit heteroscedasticity or autocorrelation. In the GLS model, temporal dependency in the residuals was corrected using an autoregressive model (corAR1), and heteroscedasticity was addressed by applying a variance function to fix variance weights. The power of a variance covariate function was selected based on the lowest AIC. The resulting model accuracy was evaluated using the adjusted coefficient of determination (adj R2) and residual standard error (RSE).

### Model projection

Climatic data forecasts were obtained from the Coupled Model Intercomparison Project Phase 6 (CMIP6) (Eyring et al., 2016). Data were also accessed via CDS website at coordinates closest to each of the ten sites (Supplementary Table S4). We used daily maximum near-surface air temperature, near-surface wind speed, and precipitation data from the climate model CMCC-ESM2 (Cherchi et al., 2019), and monthly sea surface temperature from UKESM1-0-LL (Sellar et al., 2019). All data were retrieved for historical and future projections for three main shared socio-economic pathways: SSP26, SSP45 and SSP85.

### Yearly cumulative-product survival

Future survival projections were made by applying survivability linear equations obtained in this study for mortalities observed on day 20. These equations were derived for each population using water and low tide peak temperatures from the experiments as predictors (Supplementary Table S2). Since there were no deaths in the control treatments throughout the whole experiment, the mortalities observed on day 20 were considered a delayed consequence of the 10-day heat stress event.

To adapt the daily predicted *T*_max_ data to the 10-day survival outcomes obtained in this experiment, daily *T*_max_ temperatures were averaged using a 10-day moving average algorithm. This approach enabled us to project physiological outcomes over different temporal periods and scenarios (historical, SSP26, SSP45 and SSP85). Using a 10-day average prediction showed better correlation with *in situ* measurements, improving correlations from 0.81 (daily) to 0.91 (10-day average) predictions. To account for stress accumulation in populations, a cumulative product approach was applied, generating a minimal survival rate per year.

Additionally, we simulated the yearly cumulative survival of all populations under two sets of future climate scenarios. The first set was used to evaluate the relative performance of different populations under the same environmental conditions, similar to a common-garden experiment. This approach allowed us to assess the potential of each population to cope with the same heat stress levels. Specifically, we simulated the fate of all populations under the projected climate of Viana do Castelo, the southernmost limit of the species, and presumably the most stressful location.

The second set of scenarios predicted the yearly cumulative survival of each population based on the specific climate conditions of their respective geographical locations. This allowed us to assess the likelihood of their survival in their own, realized climate.

### Population size projection

Population size at 2100 was projected by an iterative stochastic process, described by a first-order Markov chain (Caswell, 2001). This process covered the period from 2014 to 2100, corresponding to the start and ending of the CMIP6 SSP projections. The probabilistic matrices used in this projection were size-stage-classed, comprising five classes ranging from < 5 g to ≥ 190 g. Since our experimental data pertain only to fronds that fall into the first class (< 5 g), to project survivability of larger size classes, survival ratios relative to frond survival were assigned based on local demographic matrices (Supplementary Table S5). We used the Iberian matrix for the southern region (populations 1 to 4) and the Brittany matrix for the central region (populations 5 to 8) both developed by Araujo et al. (2014), and the Swedish demographic matrices from Åberg (1992a) for the northern region (populations 9 and 10).

To model the population growth, the number of individuals n(*t*+1) was calculated by multiplying each stage-class matrix probability vector (A_t_) by the yearly survival vector (S) and the previous year’s number of individuals n(*t*). The resulting stochastic matrix population model can be expressed as:

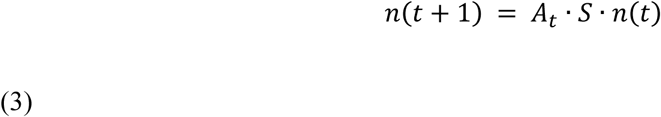

Survival (**S**) is the product of the yearly survival rate (0 to 1) with the proportional survival vector. A limiting aspect of these demographic matrices is the reduced stochastic growth rate (λ_s_) (Leslie, 1945). To balance this, we followed Åberg (1992b) approach, and scaled fertility across all classes. The scaling factor used aimed for a stable population between the years of 1850 and 2014. This was achieved by applying Equation 3 with **S** obtained from survival modelled with CMIP6 historical data from 1850-2014 (Supplementary Table S5). This can be seen as an optimistic perspective, assuming there has not been a notorious contraction in recent decades, thereby highlighting the importance of any negative outcome we might predict.

All data and code require to replicate these results are available on a Zenodo archive (Pereira et al., 2024).

## Results

### Experimental heat stress simulations

After 10 days of simulated heat stress, average growth rates tended to decrease as the temperature difference between high tides and low tides increased (Fig. 2). In other words, growth appeared to be inversely proportional to the experimental daily warming rates. In treatments with low temperature difference between high and low tide, *A. nodosum* exhibited faster growth (e.g., an average of 1.83 ± 0.88 % RGR per day in the control treatment with 8 °C temperature increase, from 20.5 °C at high tide to a maximum of 28.5 °C at low tide). Conversely, growth was inhibited in treatments with the steepest daily temperature increases (e.g., an average of –1.38 ± 1.23 % RGR per day in the treatment where temperatures increased by 25.5 °C, from 15 °C at high tide to a maximum of 40.5 °C at low tide).

**Fig. 2.**
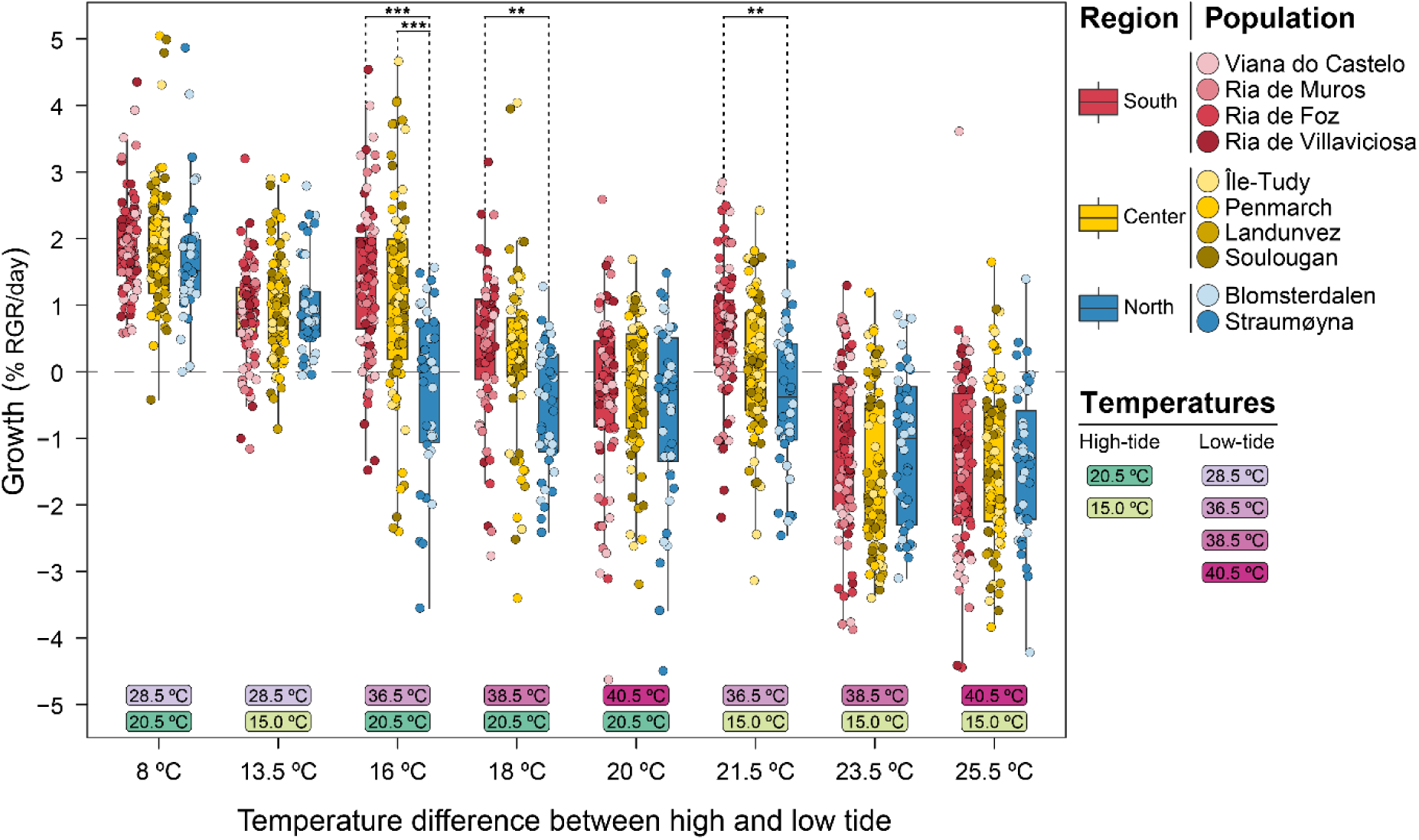
Average relative growth rate per day (% RGR day-1) (mean ± SD; n = 20 per population) after ten days of thermal stress. Experimental treatments are arranged from left to right based on the absolute temperature difference between lower temperatures at high tide and peak heating temperatures at low tide. Therefore, absolute differences ranged from 8 °C to 25.5 °C. Experimental temperatures are represented in shades of green (high-tide temperatures) and pink (low-tide peak temperatures). Points show individual growth rates, while bars show the overall interquartile distribution for each region, color-coded as blue (north populations), yellow (center populations), and red (south populations). Statistical analysis focused only on pairwise comparisons between regions within the same treatment. Significance levels: * p < .05, ** p < .01, *** p < .001.

Generally, growth rates varied inversely with the latitude of source populations. These differences were statistically significant at intermediate temperature differences of 16 °C (*p* < .001 for both), 18 °C (*p* < .02), and 21.5 °C (*p* < .001). Under these conditions, southern populations showed significantly higher growth rates compared to northern populations. At the 16 °C-increase treatment, central populations also differed significantly from their northern counterparts (*p* < .001).

By day 20 of the experiment, after the recovery period, the trend of decreasing performance from south to north persisted, particularly at intermediate temperature differences (20°C, *p* < .001 and *p* < .02; 21.5°C, *p* < .05), where differences were statistically significant (Supplementary Fig. S1). However, the inverse relationship between growth and daily warming rate, which was evident on day 10 immediately following the heat stress period, was no longer as clear. Additionally, survival rates were not evenly distributed across regions nor experimental temperatures (Fig. 3). The potential bias introduced by differential survival rates likely affected extreme temperatures and northern populations the most, followed by milder temperature increases and central and southern populations.

**Fig. 3.**
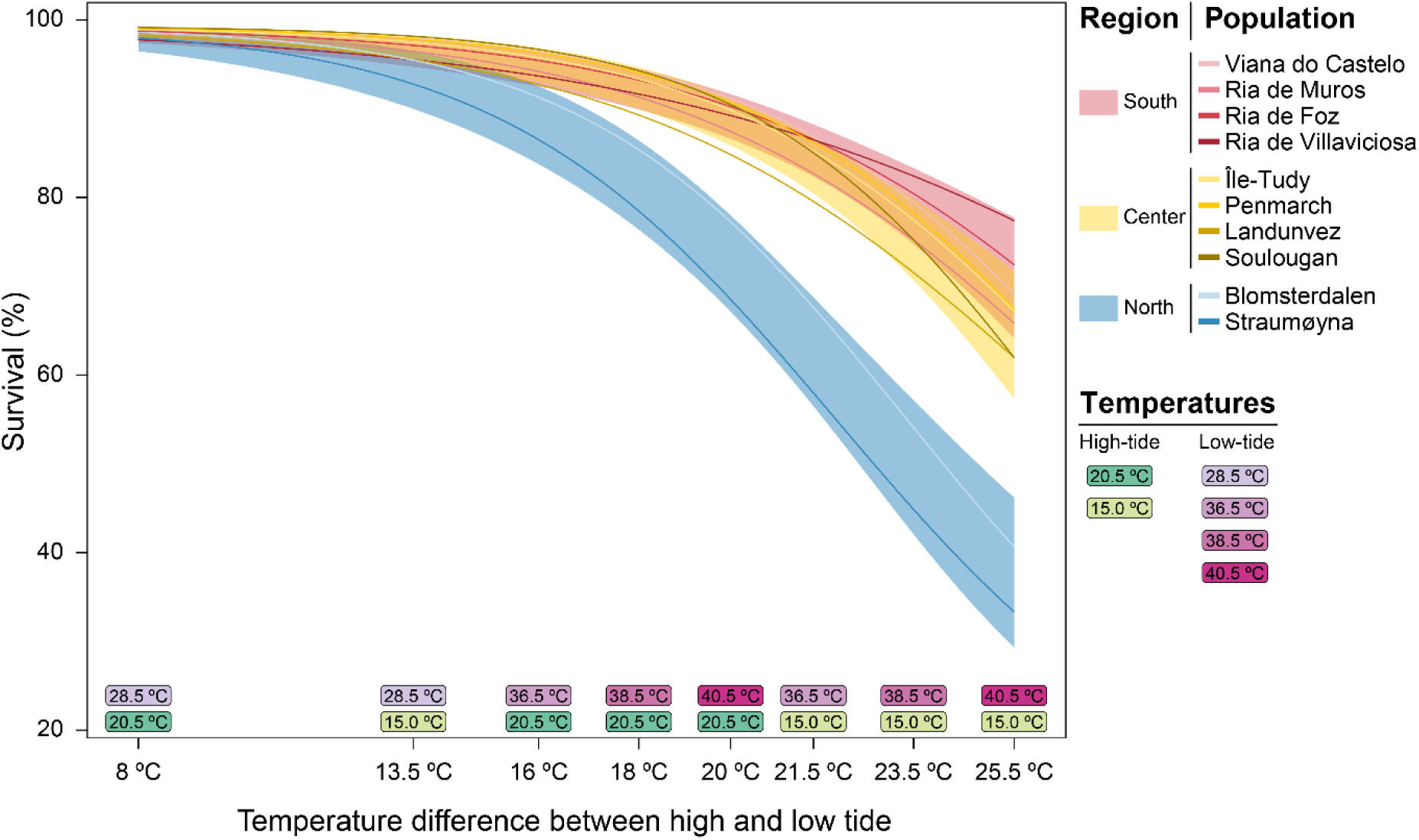
Linear model of survival rate of *A. nodosum* fronds at the end of the experiment (day 20). Treatments are arranged from left to right based on the absolute temperature difference between lower temperatures at high tide and peak heating temperatures at low tide. Treatment temperatures are represented in shades of green (high-tide temperatures) and pink (low-tide temperatures). Shaded areas show the model’s regional predictions with 95% confidence intervals, and lines represent the survival of each population. Populations from north are shown in blue, center in yellow, and south in red.

Survival rates, assessed immediately after the heat stress period on day 10 (Supplementary Fig. S2), were reduced across all treatments and did not differ significantly between regions. However, by the end of the experiment (day 20), survival rates were also inversely proportional to the absolute temperature difference between temperatures at high tide and the peak heating temperatures at low tide (Fig. 3). The highest mortality occurred with the highest temperature increases, and, consistent with growth rate trends, mortality was highest in algae from the northern populations. The lack of overlap confidence intervals in Fig. 3 (coloured shades) indicates significant differences between those populations and the center and south populations for treatments with temperature differences of >16 °C. At the highest temperature difference (25.5 °C), mortality rates were 68.9 ± 5.4% for the populations from the southern region, 56.3 ± 2.2 % for those from the central region, and 42.5 ± 2.5 % for those from the northern region (regional averages ± SD). No mortality was observed in the control treatments, regardless of the source population.

Fig. 4 shows the oxygen production and respiration of *A. nodosum* from different regions on day 10 across a range of low-high tide temperature differences. The overall trend indicates that as the low-high tide temperature difference increases, maximum net primary production (NPP_max_, upper portion of each coloured bar in Fig. 4) declines across all regions, with a more pronounced decline in the central and northern regions, suggesting greater sensitivity to heat stress.

**Fig. 4.**
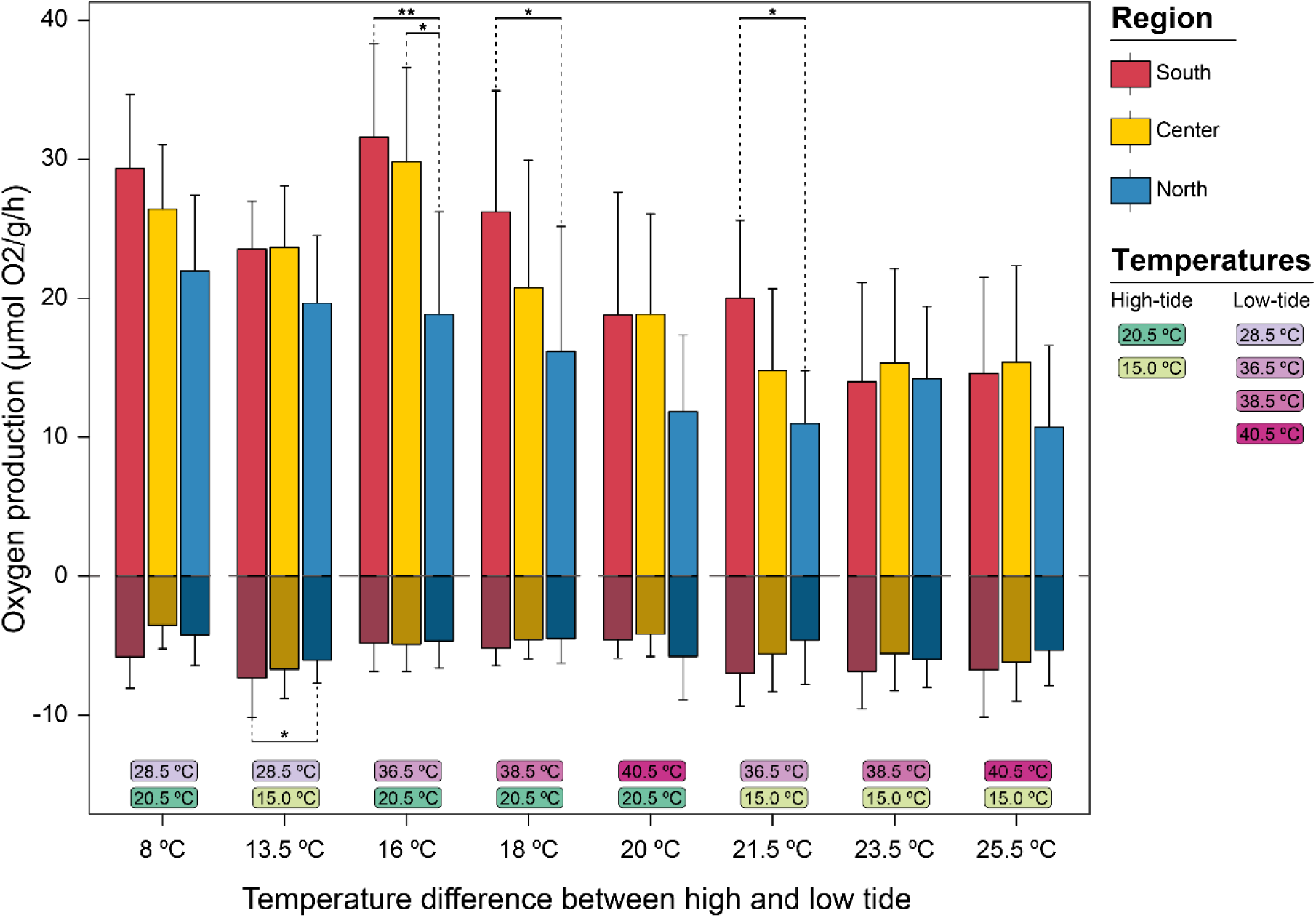
Maximum net primary production (Nppmax; µmol O2 g-1 h-1) and maximum respiration rate (Rmax; µmol O2 g-1 h-1) (mean + SD; n = 24 per region for south and center; n=12 for north) after the heatwave stress (on the day 10). Regions are color-coded as blue (north populations), yellow (center populations), and red (south populations). Nppmax is shown as positive values, while Rmax is shown as negative values (and indicates consumption). The treatments are arranged from left to right based on the absolute temperature difference between lower temperatures at high tide and peak heating temperatures at low tide. Therefore, absolute differences ranged from 8 °C to 25.5 °C. Treatment temperatures are represented in shades of green (high-tide temperatures) and pink (low-tide temperatures). Statistical analysis focused only on pairwise comparisons between regions within the same treatment. Significance levels: * p < .05, ** p < .01, *** p < .001.

Regarding regional differences, NPPmax was higher in southern populations at most temperature ranges, particularly at 16 °C (*p* < .001), 18 °C, and 21.5 °C (*p* < .05 for both), where the differences from the northern population were significant. At a temperature difference of 16 °C, algae from central populations also showed significantly higher net production compared to those from the north (*p* < .04). Maximum respiration rates (R_d_), indicated by the lower portion of each bar, showed some variability and a slight increase at the highest low-high tide temperature differences, but were generally lower for algae from the north, with statistically significant differences only in the control treatment at 13.5 °C (*p* < .05).

On day 20, NPP_max_ followed a similar but less pronounced pattern (Supplementary Fig. S3). While the northern populations continued to show a tendency for lower NPP_max_, significant differences were only found in the treatment with a warming rate of 16 °C, but only between the southern and northern populations (p < 0.05). R_d_ showed little variation and no significant regional differences in any of the treatments.

### Population forecasts

The GLS model, designed to predict *in situ* daily maximum temperatures of intertidal rock from weather data, explained 56 % of the variability observed in temperature logger data between 2021 and 2023 (Pearson’s correlation = 0.75, RMSE = 1.40 °C). The model showed that air t2m and sst had the strongest positive correlations with *in situ* temperature readings, while wind velocity and tp were negatively correlated.

Under the same climatic conditions as those in the southernmost limit of *A. nodosum* distribution (Fig. 5, left panel), the results suggest that northern populations would be severely affected. These populations show low yearly survival rates, even in the mildest scenario (SSP26), accounting 189 years (of the 250 analysed) with reduced survivability. In contrast, central and southern populations performed much better, although they experienced increased mortality under the most severe climate scenarios (SSP45 and SSP85), particularly after the 2040s.

**Fig. 5.**
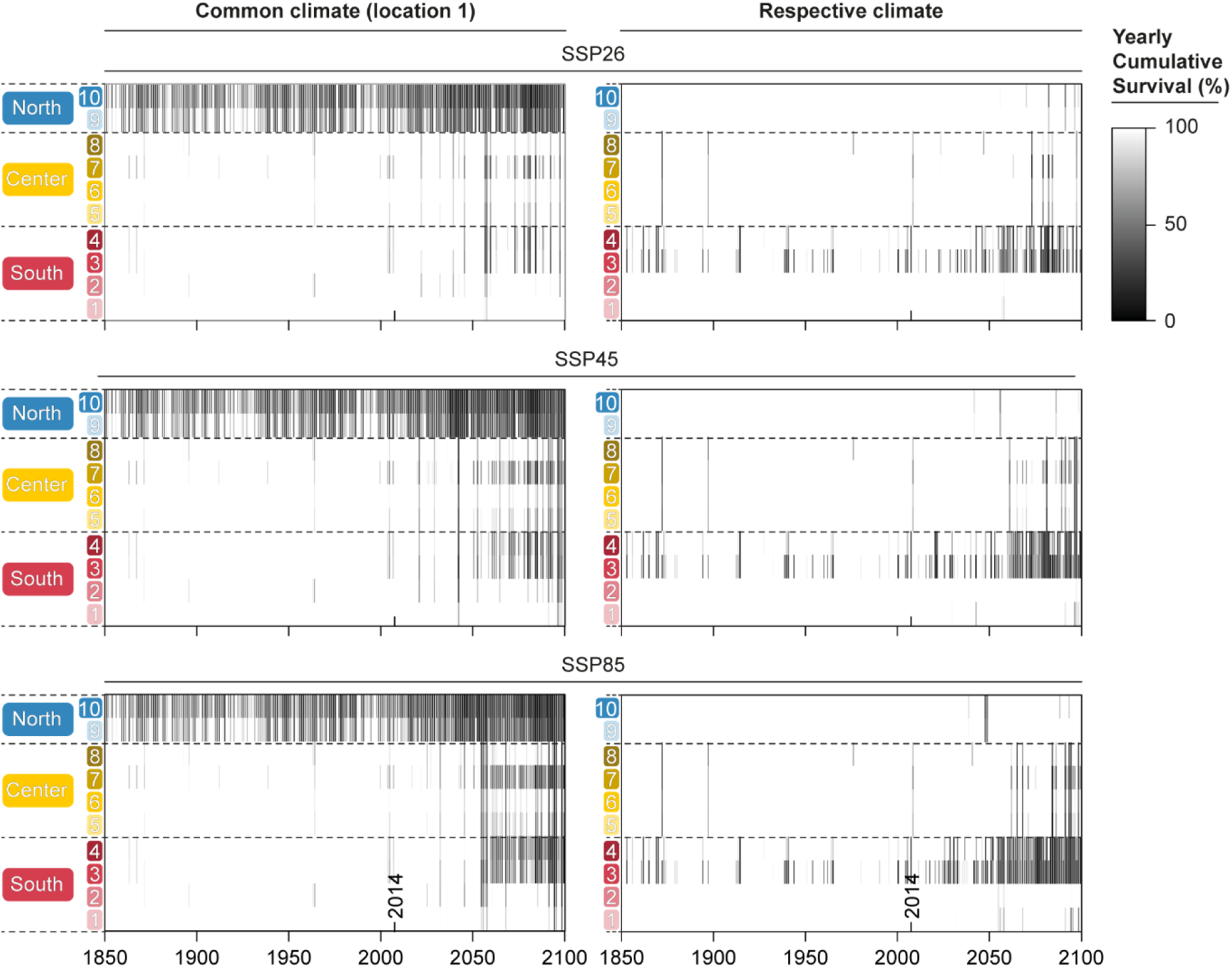
Yearly cumulative-product survival results for all ten populations from 1850 to 2100 under different SSP scenarios (SSP26, SSP45, and SSP85). The x axis represents time in yearly increments, while the vertical axis shows the regions and their respective populations. Final survival rates for each year are depicted as heatmap bars in grayscale (black represents 0% survival, and white represents 100% survival). Survival rates are reset to 100% at the beginning of each year. Plots in the left column, labelled “common climate,” show survival rates based on population-specific survivability equations applied to a single common climate (in this case, the climate experienced by the southernmost population, at Viana do Castelo, population 1, Portugal). Plots in the right column, labelled “respective climates,” indicate survival rates based on population-specific survivability equations applied to their own local climates.

On the other hand, the cumulative survival rates simulated for the specific climate conditions of each population showed a markedly different outcome (Fig. 5, right panel). Populations from northern and central Europe appeared to perform relatively well, with only a slight decline in survival rates projected by the end of the century, particularly in populations 5 and 7. A similar pattern was observed for the two southernmost populations (1 and 2), which are unlikely to experience increased heat-associated mortality over the next century. However, a different trend was seen for populations 3 and 4.

Projections in relative population sizes from 2014 to 2100 under three SSP scenarios (SSP26, SSP45, and SSP85) are shown in Fig. 6. This simulation assumes stable populations throughout the historical period of the CMIP6 hindcasts (1850 to 2014), meaning that if the frequency and intensity of heatwaves were to remain constant in the future (2014 to 2100), the populations would retain their 2014 sizes.

**Fig. 6.**
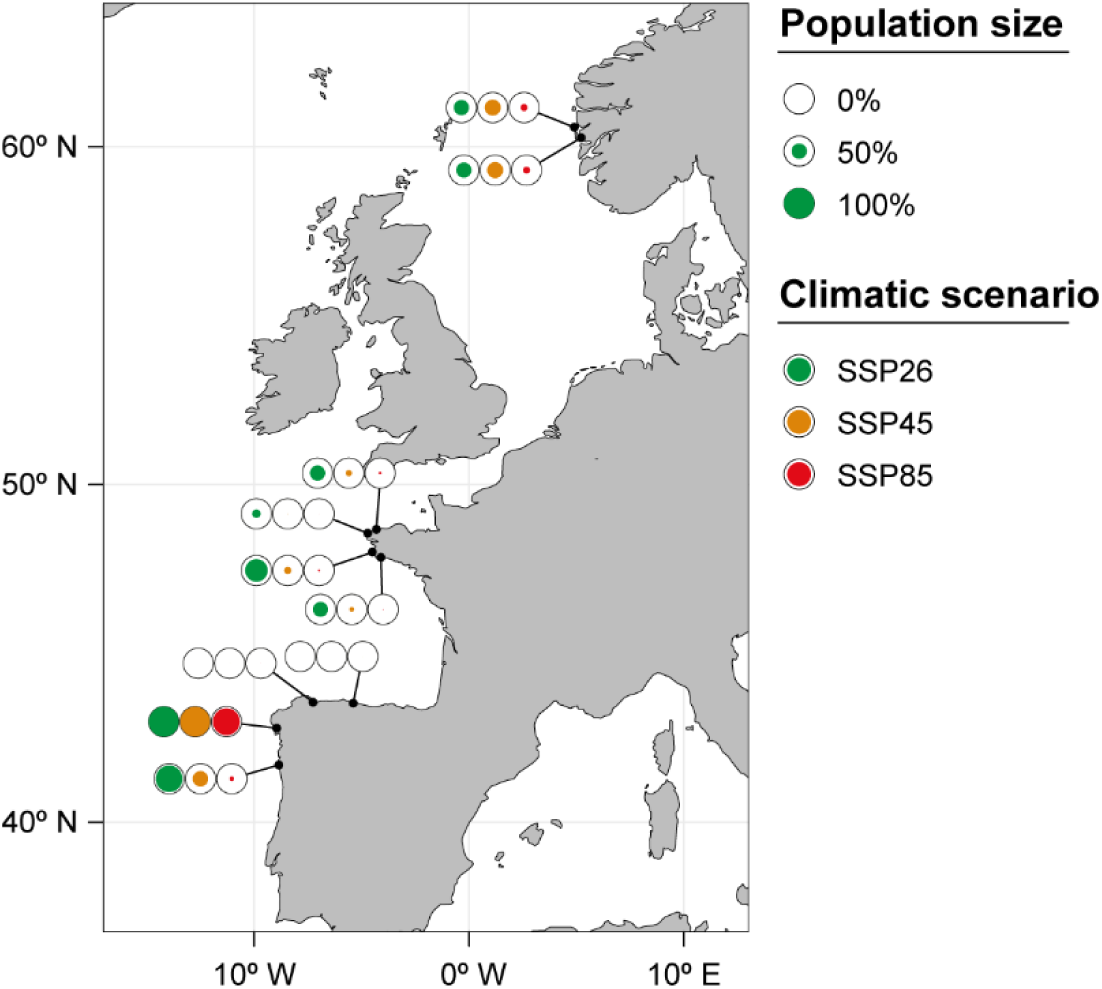
Projected population sizes for all ten populations by the end of 2100 under different SSP scenarios (SSP26, SSP45, and SSP85). Black circumferences represent the population size in 2014, while coloured filled circles indicate the expected population size in 2100, with the size of each circle proportional to the population size. Filled circles smaller than the black circumference indicate population contraction, while empty circumferences represent population extinction. Each projection considers respective climate conditions (based on CMIP6 climatic data), yearly survival rates, and regional demographic matrices. The map was created in R (R Development Core Team, 2023) using the Rnaturalearth library v1.0.0.

Firstly, it is observed that, for some locations, population sizes are expected to remain mostly stable. The most positive forecast is for the population of 2 (Ría de Muros) in NE Iberia, where the resilience of local individuals, combined with projected local warming, suggests that the population will likely maintain a relatively good demographic condition across all SSP scenarios. Relatively high population sizes are also expected for other locations, such as population 1 (Viana do Castelo) in NE Iberia and populations 5 (Île-Tudy), 6 (Penmarch), and 8 (Soulougan) in Brittany, but only under the mildest warming scenario (SSP26). Significant population size reductions are predicted under the two more severe SSP scenarios in all these locations, suggesting vulnerability to extreme climate change conditions.

Secondly, the two northernmost populations, 9 and 10 (Blomsterdalen and Straumøyna), exhibit intermediate and similar responses under the SSP26 and SSP45 scenarios but are also projected to be strongly affected under the SSP85 climate scenario.

Thirdly, there are populations projected to face severe reductions or even extinction, irrespective of the climate scenario, emphasizing their heightened susceptibility to climate change. These include populations 3 (Ría de Foz) and 4 (Ría de Villaviciosa) in northern Iberia and population 7 (Landunvez) in northern Brittany.

While a few populations may demonstrate resilience and maintain stable sizes under moderate climate change scenarios, others, particularly those in northern Iberia and Brittany, are at risk of significant decline or extinction.

## Discussion

This study aimed to investigate the physiological responses of the brown seaweed *Ascophyllum nodosum* to simulated intertidal heat stress across its distributional range. By replicating realistic intertidal thermal conditions, with precise control over tides, light, and daily temperatures, we were able to (i) assess the significance of upwelling events at the southern limit of the species’ distribution; (ii) identify thermal patterns with physiological stress; (iii) demonstrate population-specific tolerance to thermal stress; and (iv) produce long-term population forecasts throughout the European range of this species.

### The importance of cold, upwelled waters

Our data suggest that *A. nodosum* does not directly benefit from the relatively cold waters associated with upwelling pulses at its southern distribution limit. Given its Arctic-Boreal distribution, to approximately 42° latitude in northern Portugal (Pereira et al., 2020), one could expect this species to be limited by warm waters (Lüning, 1990). However, our findings indicate that *A. nodosum* performs better at all treatments with water temperatures of 20.5 °C compared to 15 °C, suggesting that, unlike other cold-water species (Wilson et al., 2015), it may grow better during upwelling relaxation. A similar conclusion was reached by Keser et al. (2005), who studied the growth of *A. nodosum* in the field at the southern distribution limit of the species in Connecticut, USA. More recently, it was demonstrated in the lab that growth increased with temperatures up to ∼25 °C, although these experiments did not include any period of aerial exposure (Hernández et al., 2023; Wilson et al., 2015)

In *Ascophyllum nodosum*, as anticipated from its growth rates, productivity has also been shown to improve with increasing water temperatures up to approximately 22 °C under natural field conditions (Chock & Mathieson, 1979). Laboratory studies by Piñeiro-Corbeira et al. (2018) have demonstrated that mild increases in water temperature can enhance photosynthesis and respiration in other Fucales, although this positive effect is quickly reversed at critical temperatures between 21°C and 22°C. The warmest seawater temperature in our experiments (20.5°C) remained below these thresholds, which likely accounts for the lack of observed detrimental effects of warm water on net primary productivity.

Overall, our findings may explain why the southern limit of *A. nodosum* has remained stable in northern Portugal, where sea temperatures rarely exceed 22 °C (Seabra et al., 2011), while other cold-water species have experienced significant range contraction northwards (Casado-Amezúa et al., 2019; Lima et al., 2006). This realization raises the question of what actually sets its distribution limit. One contributing factor may be the scarcity of suitable habitat. *A. nodosum* forms dense populations only on wave-protected to moderately protected rocky shores (Scrosati and Heaven, 2007). South of Viana do Castelo, in northern Portugal, such protected shores are rare, and the few that do exist are located deep inside river mouths or coastal lagoons. Our results, however, also point to another potential limiting factor: thermal stress resulting from the temperature difference between each consecutive high and low tide.

### Thermal stress response from high to low tides

Given their relatively high position on the shore, many intertidal fucoids have developed physiological adaptations to cope with stressors such as increased ultraviolet radiation, elevated temperatures, and desiccation during low tide. In fact, the upper distribution limits of intertidal algae are believed to be primarily determined by their capacity to withstand and recover from low-tide emersion (Schonbeck and Norton, 1978; Wiltens et al., 1978; Dring & Brown, 1982). However, not all the effects of low tide are detrimental. As documented by Dring and Brown (1982), Nitschke et al. (2010), and Ryzhik et al. (2018) for various brown algae species, increased light exposure during emersion can enhance photosynthetic activity, at least during the initial stages of exposure.

Although important, realistic experiments involving low tides remain rare and, to our knowledge, have never been conducted with *A. nodosum*. Consequently, directly comparable data from low-tide assessments are lacking. Nonetheless, our growth control results align with previously reported optimal daily growth rates of 2% to 3% (Fortes, 1980). The inhibited growth observed in the treatments featuring higher low-tide temperatures was likely due to damage to the photosynthetic system caused by stress factors. This is supported by the observed decrease in oxygen production, indicating limited carbon fixation, lower ATP production, and, consequently, reduced growth (Takahashi & Murata, 2008). Additionally, this inhibition is unlikely to be due to increased metabolic costs, as this would typically result in higher respiration rates (Hurd et al., 2014), which were not observed. Instead, the results suggest an overall compromise of the photosynthetic apparatus (Eggert, 2012).

Experiments with other species also suggest that high temperatures during emersion may impair growth more severely than when seaweeds are submerged. Román et al. (2020) demonstrated that even mild low-tide heat stress can reduce growth in macroalgae far more than marine heatwaves. Similarly, experiments on *Fucus serratus* found that low-tide treatments at 34°C had a greater impact on growth than hot water treatments at 22°C (Martinez et al., 2012).

Although our results indicate a general decrease in performance with increasing low-tide thermal stress, this relationship was not entirely linear. The treatments in which the algae showed the poorest performance — whether measured by survivability, growth, or respiration — were not necessarily those with the highest temperature peaks. Instead, the lowest performance occurred in treatments where the algae experienced the most temperature increase, from the time when the tide receded until the temperature peak at low tide. Interestingly, similar research on a cold-adapted intertidal gastropod (*Patella vulgata*) by Seabra et al. (2016) found that heat stress over a 30-day period was directly linked to elevated water temperatures, with high air temperatures causing stress only when water temperatures were also high. Hence, one cannot assume that being repeatedly exposed to upwelling relaxation has no deleterious consequences for *A. nodosum* since longer-term effects (i.e.,> 20 days) in performance and survival remain to be tested. However, because realistic simulations of the entire tidal cycle are rarely done, connecting these findings to broader ecological implications remains challenging, and further research is needed to determine whether these patterns can be generalized to other species and geographic locations.

Nonetheless, our study underscores the importance of designing laboratory experiments that accurately simulate the complex thermal environments experienced by intertidal species. Undeniably, laboratory simulation deemed realistic can be challenging, as these require fine control of both marine and terrestrial conditions. Moreover, the technical challenges extend beyond the experimental setup to the data collection phase, as it is crucial to gather long-term temperature data across both high and low tides at the scale of the studied organisms (Lima et al., 2009). Given these complexities, it is unsurprising that many studies focus only on simulating high tide conditions, overlooking the thermal stress experienced during low tide. This approach carries the risk of overestimating the ecological niche of intertidal species. In the case of *A. nodosum*, this may help explain the large-scale discrepancies between its predicted and actual observed range reported by Hernández et al. (2023).

### Thermal sensitivity across the species distribution

Common-garden experiments, in which different populations of a species are exposed to the same environmental conditions, are often used to determine which populations are better adapted to specific climatic conditions (Schwinning et al., 2022). In this context, our results were quite clear: across all evaluated parameters — survival, growth, and oxygen production — a consistent performance gradient emerged from northern populations performing the poorest and southern populations exhibiting the highest performance. Differences were most pronounced in the mid-range treatments, likely because, at the most benign levels, the stress was not severe enough to elicit strong responses, while at higher stress levels, organisms were already experiencing damage, regardless of their population origin (i.e., photosystem damage, oxidative stress, Allakhverdiev et al. 2008, Takahashi & Murata, 2008).

While regional differences were found in both growth and oxygen production, the most significant differences in thermal sensitivity were registered in survival rates, in agreement with other studies that compared survival and growth performance (Bennett et al., 2015; Mohring et al., 2014). Interestingly, in our experiment, most mortality did not occur immediately following heat stress but emerged later in a delayed manner. This is also in agreement with previous reports by Edwards (2004), Raymond et al. (2022), and Speare et al. (2022).

Our results have some inherent limitations that should be acknowledged. First, they do not provide insight into the mechanisms underlying the latitudinal cline in performance (King et al., 2017). Thus, we cannot distinguish between local adaptation (where natural selection causes changes in allele frequencies, leading to a shift towards local optimal traits) and phenotypic plasticity (where individuals adjust their phenotype in response to local conditions through changes in gene expression), although, within the scope of this study, this distinction is not critical. Secondly, our laboratory trials, despite efforts to closely simulate the thermal environment of the intertidal zone, cannot fully replicate the complex natural conditions these organisms face in the wild. Field thermal stress depends on the resilience, recovery, and recolonization traits of the affected species, along with interactions with other stressors such as eutrophication, altered currents, salinity, or modified grazing pressure (Straub et al., 2019), none of which were simulated in this study.

### Population hindcasts, forecasts, and extinction risk

The modelled survival of populations under a hypothetical common climate reflects the results of the common-garden experiment. It indicates that northern populations, if exposed to the warmer temperatures typical of Viana do Castelo, Portugal, would have experienced frequent mortality events in the past and would likely continue to do so in the future. In contrast, central and southern populations would have remained largely unaffected in the past and would only begin to show differences starting in the mid-2040s. This suggests that while central and southern populations are adapted to the warmer conditions they have been experiencing, northern populations would struggle if exposed to those same temperatures. Such heterogeneous responses are expected in highly isolated populations. *A. nodosum* is well-known for its limited dispersal (Badas et al., 1990), and genetic evidence indicates that southern populations may have drifted apart due to low gene flow (Olsen et al., 2010).

These patterns were substantially different when survival was simulated under each population’s respective climate. Northern and central populations appear to have been well-adapted to the climates they have recently experienced. Looking ahead, northern populations show moderate declines under the milder scenarios, with slightly better performance in SSP45 than in SSP26. This aligns with other studies suggesting that mild warming may expand the ecological niche for *A. nodosum* at higher latitudes (Jueterbock et al., 2013; Westmeijer et al., 2019). Central populations, on the other hand, show moderate contractions under the SSP26 scenario but experience a sharp collapse under harsher scenarios, consistent with the partial extinction projected for Brittany by Jueterbock et al. (2013).

In the southern region, there was greater variability among populations. While population 1 is predicted to experience moderate losses under the most severe conditions, population 2 is expected to perform the best across all SSP scenarios. This may be a consequence of its geographical setting (i.e., inside a large Ría, a high-relief drowned river valley), and hints at the importance of such enclosed environments, which may buffer coastal water temperatures (Sousa et al., 2020). On the other hand, the two remaining southern populations, 3 (Ría de Foz) and 4 (Ría de Villaviciosa), seem to have experienced frequent mortality events in the past, and these are expected to worsen rapidly within this century. This result matches recent biogeographic changes in this region, where several fucoid species have been retreating westward to the relatively cooler northwest corner of the Iberian Peninsula (Casado-Amezúa et al., 2019; Fernández, 2011). For instance, the distributions of *Fucus serratus* and *Himanthalia elongata* have contracted, respectively, by 197 km and by 330 km since the end of the 19th century in northern Spain (Duarte et al., 2013). The distribution of *A. nodosum* has similarly shifted westward. Our survey found the easternmost population of *A. nodosum*, consisting of only five individuals, in Niembru, northern Spain — approximately 218 km from Pedernales, the distribution limit documented by Fischer-Piette in the 1950s (Fischer-Piette, 1955). Furthermore, populations 3 and 4 are currently inhabited by smaller individuals that occupy only a few tens of square meters, in contrast to the other populations, which cover hundreds of square meters and consist of much larger (and seemingly healthier) individuals (L. Pereira, pers. observation). This scenario contradicts the general forecasts made by Jueterbock et al. (2013) and Westmeijer et al. (2019) of an overall northward contraction of the species’ range at its southern limit. However, their correlative approach did not account for the importance of local ecotypes, which may explain these discrepancies.

This modelling exercise is based on several assumptions and should, therefore, be interpreted with caution. First, the model relies solely on the mortality rates observed in this study and does not account for any sub-lethal effects of heat stress. Second, converting CMIP6 projections to actual maximum temperatures in the intertidal zone relied on a correlational relationship between meteorological data and *in situ* observations, with inherent uncertainties. Despite these potential limitations, the model performed extremely well during the hindcast period (1850–2014) when we simulated the yearly cumulative survival of each population under its respective observed climate. This provided an opportunity for model validation as it showed that, during the 20th century, no populations appeared to be severely impacted by heat stress except for populations 3 and 4, precisely in the region where distribution retractions have been observed.

Even if we take a cautious approach and refrain from interpreting the modeling results in absolute terms, they still offer valuable insights. They illustrate how varying sensitivities of populations across a species’ range, combined with region-specific climate forecasts, can create a geographic mosaic of outcomes. Range contractions may occur at the warm edge of the distribution, where populations, though more resilient to thermal stress, can still be overwhelmed by the rapid pace of warming. In contrast, despite being more sensitive to heat stress, northern populations may be less negatively affected by future climate.

In conclusion, this study underscores the importance of conducting realistic experiments when evaluating a species’ thermal tolerance. Our findings indicate that *Ascophyllum nodosum* populations exhibit a fairly latitudinal cline in sensitivity to heat stress. By integrating empirical physiological observations with state-of-the-art climate projections, we were able to significantly refine forecasts at the population level. Our results show that future climate conditions can, in some situations, outweigh the differences in thermal stress resilience, potentially leading to uneven impacts on various populations along the species’ current distribution.

## Acknowledgments

This work was supported by Fundação para a Ciência e Tecnologia (FCT) through the projects ThermalBuffer (POCI-01-0145-FEDER-31088) and OceanLog (PTDC/BIA-BMA/4848/202), Blue Growth program – EEA grants BLUEFORESTING (PT-INNOVATION-0077), EU H2020 FutureMARES (contract no. 869300), and CESAM (UIDP/50017/2020 + UIDB/50017/2020 + LA/P/0094/2020), funded by national funds (OE) through FCT/MCTES. LFP received support from a FCT PhD grant (SFRH/BD/136924/2018). CM was supported through the abovementioned OceanLog and Thermalbuffe, as well as the EU H2020 project ANERIS (no. 101094924). FPL, RS, RdS and SF received support from FCT through CEECInd contracts (10.54499/CEECIND/03185/2018/CP1546/CT0004, 10.54499/CEECIND/01424/2017/CP1423/CP1645/CT0006, 10.54499/2021.03268.CEECIND/CP1668/CT0001, 10.54499/2021.02653.CEECIND/CP1659/CT0012). We want to thank Consolación Fernández for directions to populations of *A. nodosum*, Kjersti Sjøtun for her support in Norway, Hugo Meyer and Óscar Gómez for their help with the lab experiments, and Per Åberg and Rita Araújo for their support and sharing of unpublished demographic matrixial data.

## Annexes

**Table. S1.**
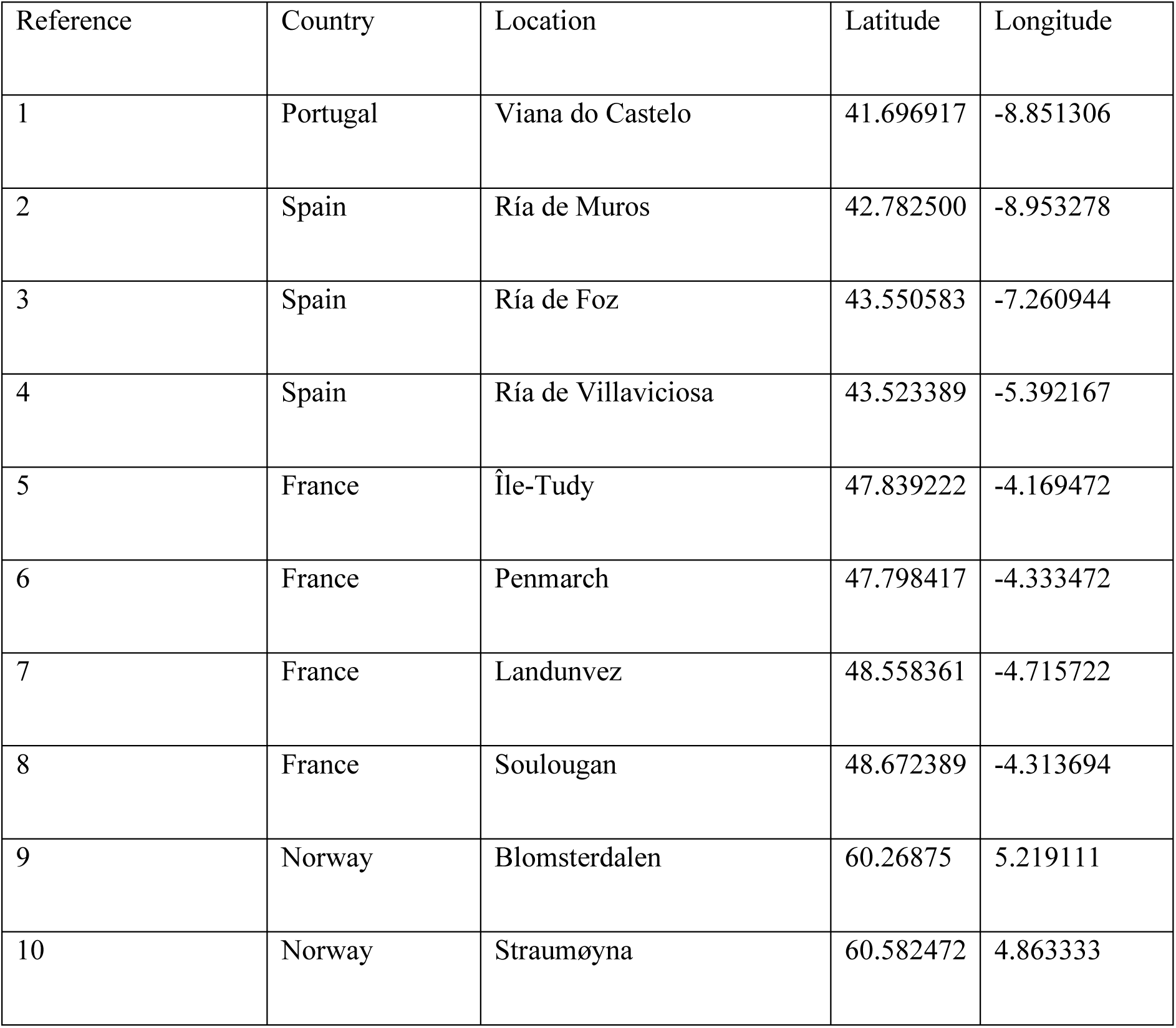
Overall coordinates for the sampled populations.

| Reference | Country | Location | Latitude | Longitude |
| --- | --- | --- | --- | --- |
| 1 | Portugal | Viana do Castelo | 41.696917 | -8.851306 |
| 2 | Spain | Ría de Muros | 42.782500 | -8.953278 |
| 3 | Spain | Ría de Foz | 43.550583 | -7.260944 |
| 4 | Spain | Ría de Villaviciosa | 43.523389 | -5.392167 |
| 5 | France | Île-Tudy | 47.839222 | -4.169472 |
| 6 | France | Penmarch | 47.798417 | -4.333472 |
| 7 | France | Landunvez | 48.558361 | -4.715722 |
| 8 | France | Soulougan | 48.672389 | -4.313694 |
| 9 | Norway | Blomsterdalen | 60.26875 | 5.219111 |
| 10 | Norway | Straumøyna | 60.582472 | 4.863333 |

**Table. S2.** Survival rate equation constants, per population. Equations followed the form “Survival rate = β_0_ + β_1_ LPT + β_2_ SST + β_3_ Tank_R_” where LPT represents low tide peak temperature, SST represents water temperature (SST), and Tank_R_ represents cross random effect of experimental tank. Models selected by their AIC.

| Reference | Location | $\beta_0$ | $\beta_1$ | $\beta_2$ | $\beta_3$ | AIC |
| --- | --- | --- | --- | --- | --- | --- |
| 1 | Viana do Castelo | 16.731 | -0.3975 | 1.6141 | 0.7429 | 99.50279 |
| 2 | Ría de Muros | 19.3459 | -0.4684 | 1.2969 | 1.029 | 113.4228 |
| 3 | Ría de Foz | 19.276 | -0.4588 | 1.0057 | 0.6357 | 101.4419 |
| 4 | Ría de Villaviciosa | 12.0576 | -0.2702 | 0.5617 | 0.5105 | 109.5692 |
| 5 | Île-Tudy | 20.2749 | -0.4902 | 1.1389 | 0.5104 | 104.0603 |
| 6 | Penmarch | 18.6274 | -0.4491 | 1.2798 | 0 | 101.2708 |
| 7 | Landunvez | 21.4076 | -0.5253 | 1.5058 | 0 | 102.4099 |
| 8 | Soulougan | 14.1529 | -0.3434 | 0.9252 | 0 | 129.3407 |
| 9 | Blomsterdalen | 13.028 | -0.339 | 1.753 | 0.102 | 145.6631 |
| 10 | Straumøyna | 12.1336 | -0.3262 | 1.6163 | 0 | 160.0565 |

**Table. S3.**
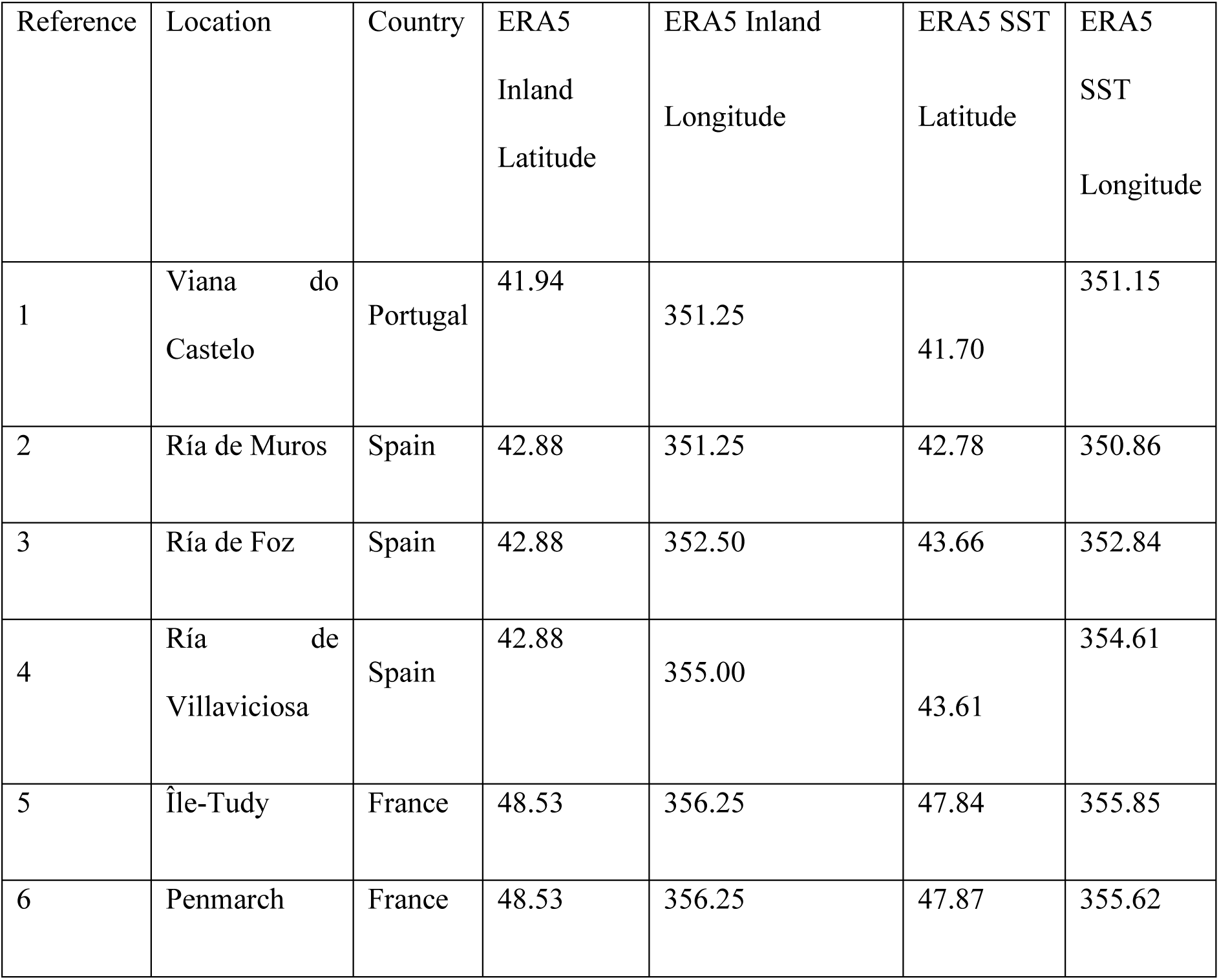
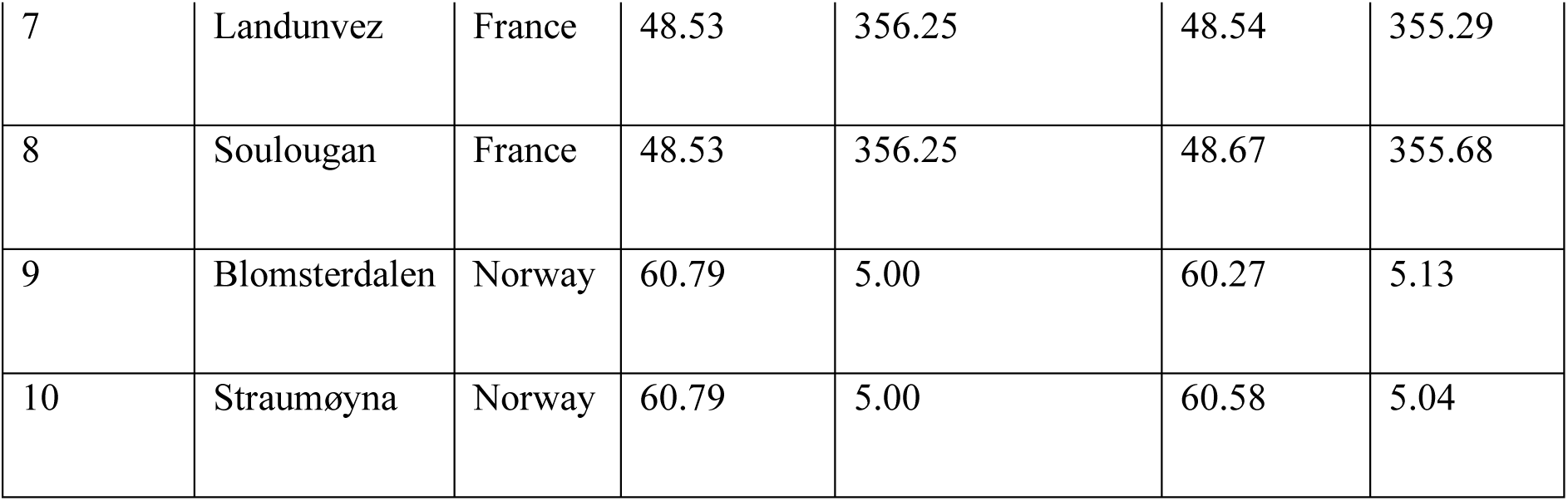
Coordinates sampled on the ERA5 dataset. Data was accessed in January 2024.

**Table. S4.**
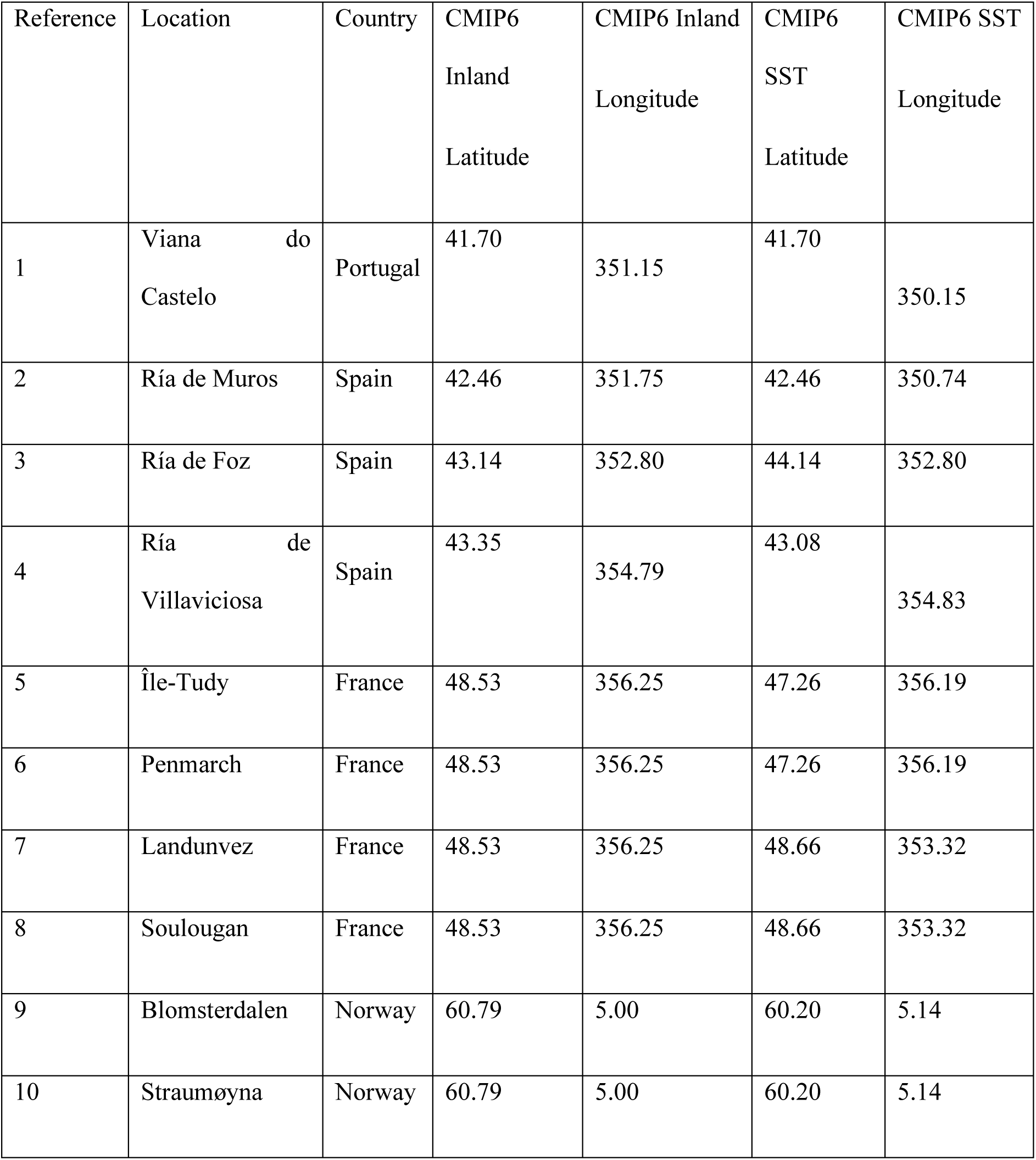
Coordinates sample on the CMIP6 dataset. Data was accessed in January 2024.

**Table. S5.** Survival ratio for the five size classes relative to size class 1, and fertility inflation ratios for all populations. Survival ratio values are organized by size class (1 to 5) and represent the natural survivability of each size class compared to size class 1. The equation used to determine a size class’s survival is: Survival rate = 1 – ((1 – Size class 1 survival rate / respective size class ratio). Survival ratios are derived from demographic matrixes obtained from Åberg (1992a) and Araujo et al. (2014). Fertility inflation was calculated based on the population’s requirement to maintain the same size in 2014 it had in 1850, using a stochastic approach with historical CMIP6 data (1850 to 2014).

| Reference | Location | Survival ratios | Fertility inflation |
| --- | --- | --- | --- |
| 1 | Viana do Castelo | 1, 1.77, 2.19, 2.25, 2.44, 1.84 | 1.84 |
| 2 | Ría de Muros |  | 1.84 |
| 3 | Ría de Foz |  | 3.84 |
| 4 | Ría de Villaviciosa |  | 2.41 |
| 5 | Île-Tudy | 1, 2, 3.25, 3.31, 3.30, 3.1 | 12.66 |
| 6 | Penmarch |  | 12.59 |
| 7 | Landunvez |  | 12.82 |
| 8 | Soulougan |  | 13.05 |
| 9 | Blomsterdalen | 1, 1.06, 1.11, 1.13, 1.15 | 1.376 |
| 10 | Straumøyna |  | 1.376 |

**Table. S6.** Summary of the results from generalized linear mixed models (GLMMs) testing the effects of low-tide temperatures (Air), hight-tide temperatures (Water), Region, Population as a nested random effect within Region (Population (Region)), Tank as cross-random effect, and their interactions for best model in each parameter. Significant effects (p < 0.05) are indicated in bold, while * denotes marginal significance. The models presented cover growth, net oxygen production and respiration, after the heat stress and the recovery periods.

| Variable | After heat stress |  |  | After recovery |  |  |
| --- | --- | --- | --- | --- | --- | --- |
| *Growth* |  |  |  |  |  |  |
| Best model | A x R + W x R + P(R) + T |  |  | A x W + A x R + P(R) + T |  |  |
| Fixed effects | $\chi^2$ | Df | p-value | $\chi^2$ | Df | p-value |
| Air (A) | 112.92 | 3 | **<0.001** | 149.05 | 3 | **0.002** |
| Water (W) | 37.07 | 1 | **<0.001** | 185.34 | 1 | **<0.001** |
| Region (R) | 19.23 | 2 | **<0.001** | 37.70 | 2 | **<0.001** |
| A x W | - |  |  | 90.90 | 3 | 0.076* |
| A x R | 50.94 | 6 | **<0.001** | 13.06 | 6 | **0.021** |
| W x R | 21.28 | 4 | **<0.001** | - |  |  |
| A x W x R | - |  |  | - |  |  |
| Random effects | Variance |  | Std. Dev. | Variance |  | Std. Dev. |
| Tank (T) | 0.102 |  | 0.320 | 0.1312 |  | 0.044 |
| Population (Region) (P(R)) | 0.001 |  | 0.001 | 0 |  | 0 |
| Residual | 1.179 |  | 1.086 | 0.925 |  | 0.9618 |
| *Net primary production* |  |  |  |  |  |  |
| Best model | A x W x R + P(R) + T |  |  | A x W x R + P(R) + T |  |  |
| Fixed effects | $\chi^2$ | Df | p-value | $\chi^2$ | Df | p-value |
| Air (A) | 76.950 | 3 | **<0.001** | 7.53 | 3 | 0.057* |
| Water (W) | 65.170 | 1 | **<0.001** | 77.53 | 1 | **<0.001** |
| Region (R) | 16.300 | 2 | **<0.001** | 10.42 | 2 | **0.005** |
| A x W | 20.590 | 3 | **<0.001** | 0.29 | 3 | 0.962 |
| A x R | 14.530 | 6 | **0.024** | 7.69 | 6 | 0.262 |
| W x R | 7.910 | 2 | **0.019** | 12.24 | 2 | **0.002** |
| A x W x R | 12.290 | 6 | 0.055* | 1.14 | 6 | 0.980 |
| Random effects | Variance |  | Std. Dev. | Variance |  | Std. Dev. |
| Tank (T) | 1.27 |  | 1.128 | 1.056 |  | 1.028 |
| Population (Region) (P(R)) | 3.25 |  | 1.803 | 3.450 |  | 3.450 |
| Residual | 42.13 |  | 6.491 | 33.637 |  | 5.800 |
| *Respiration* |  |  |  |  |  |  |
| Best model | A x W x R + P(R) + T |  |  | A x W x R + P(R) + T |  |  |
| Fixed effects | $\chi^2$ | Df | p-value | $\chi^2$ | Df | p-value |
| Air (A) | 6.49 | 3 | <0.089* | 3.09 | 3 | 0.377 |
| Water (W) | 53.83 | 1 | **<0.001** | 9.80 | 1 | **<0.001** |
| Region (R) | 24.39 | 2 | **<0.001** | 9.49 | 2 | **0.009** |
| A x W | 11.84 | 3 | **<0.001** | 3.05 | 3 | 0.383 |
| A x R | 8.27 | 6 | 0.22 | 5.53 | 6 | 0.478 |
| W x R | 6.10 | 2 | **0.047** | 2.9 | 2 | 0.234 |
| A x W x R | 3.22 | 6 | 0.78 | 4.93 | 6 | 0.552 |
| Random effects | Variance |  | Std. Dev. | Variance |  | Std. Dev. |
| Tank (T) | 0.884 |  | 0.940 | 0.001 |  | 0.001 |
| Population (Region) (P(R)) | 0 |  | 0 | 0 |  | 0 |
| Residual | 8.052 |  | 2.837 | 7.012 |  | 2.650 |

**Fig. S1.**
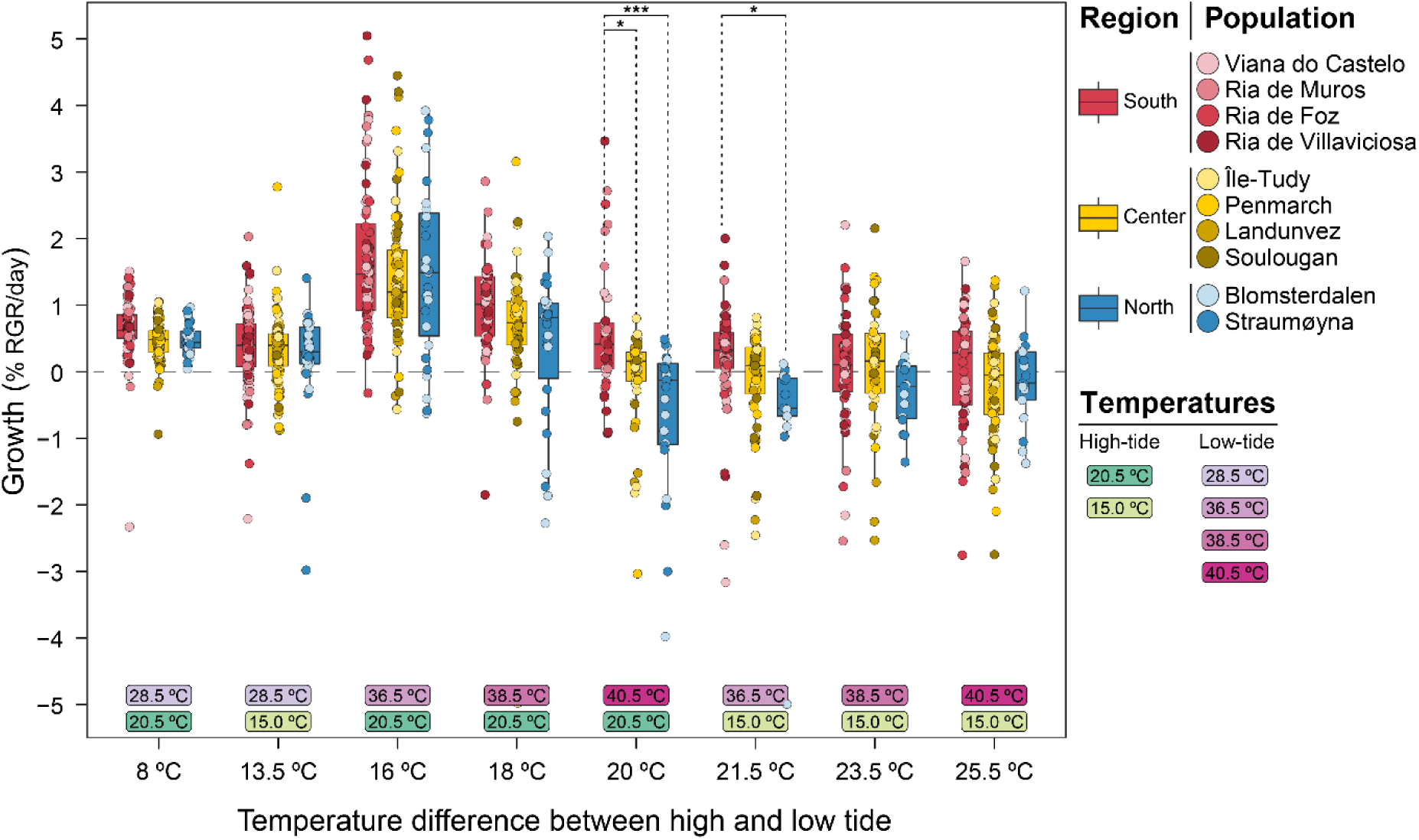
Average relative growth rate per day (% RGR day-1) (mean ± SD; n = 20 per population) after the recovery period (on the day 20). The treatments are arranged from left to right based on the absolute temperature difference between lower temperatures at high tide and peak heating temperatures at low tide. Treatment temperatures are represented in shades of green (high-tide temperatures) and pink (low-tide temperatures). Points show individual growth rates, while bars show the overall interquartile distribution for each region, color-coded as blue (north populations), yellow (center populations), and red (south populations). Statistical analysis focused only on pairwise comparisons between regions within the same treatment. Significance levels: * p < 0.05, ** p < 0.01, *** p < 0.001.

**Fig. S2.**
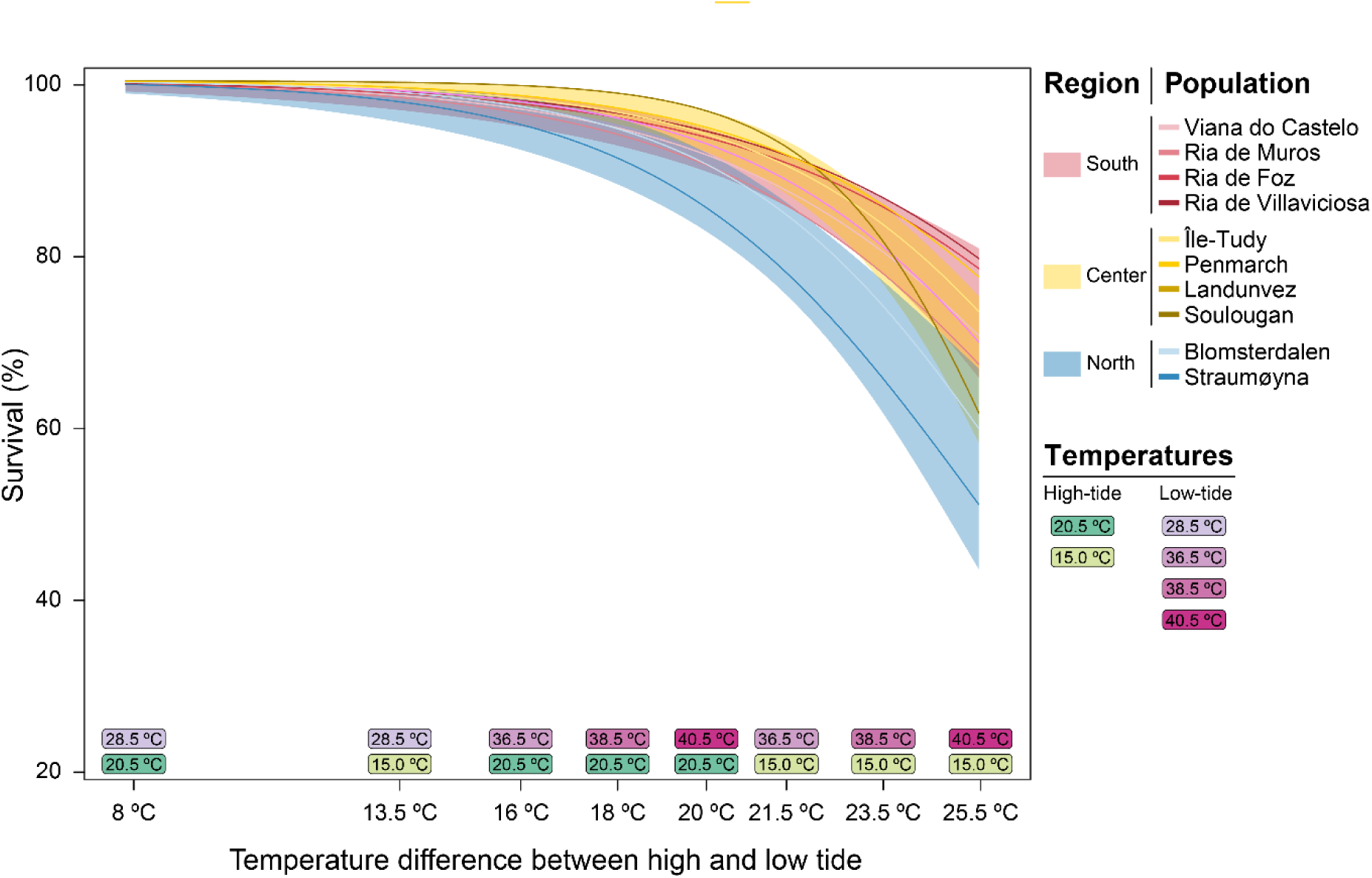
Linear model of survival rate of A. nodosum fronds after heat stress events (on the day 10). Treatments are arranged from left to right based on the absolute temperature difference between lower temperatures at high tide and peak heating temperatures at low tide. Treatment temperatures are represented in shades of green (high-tide temperatures) and pink (low-tide temperatures). Shaded areas show model’s regional predictions with 95% confidence intervals and lines represent the survival of each population. Populations from Norway are shown in blue, France in yellow, and Iberian Peninsula in red.

**Fig. S3.**
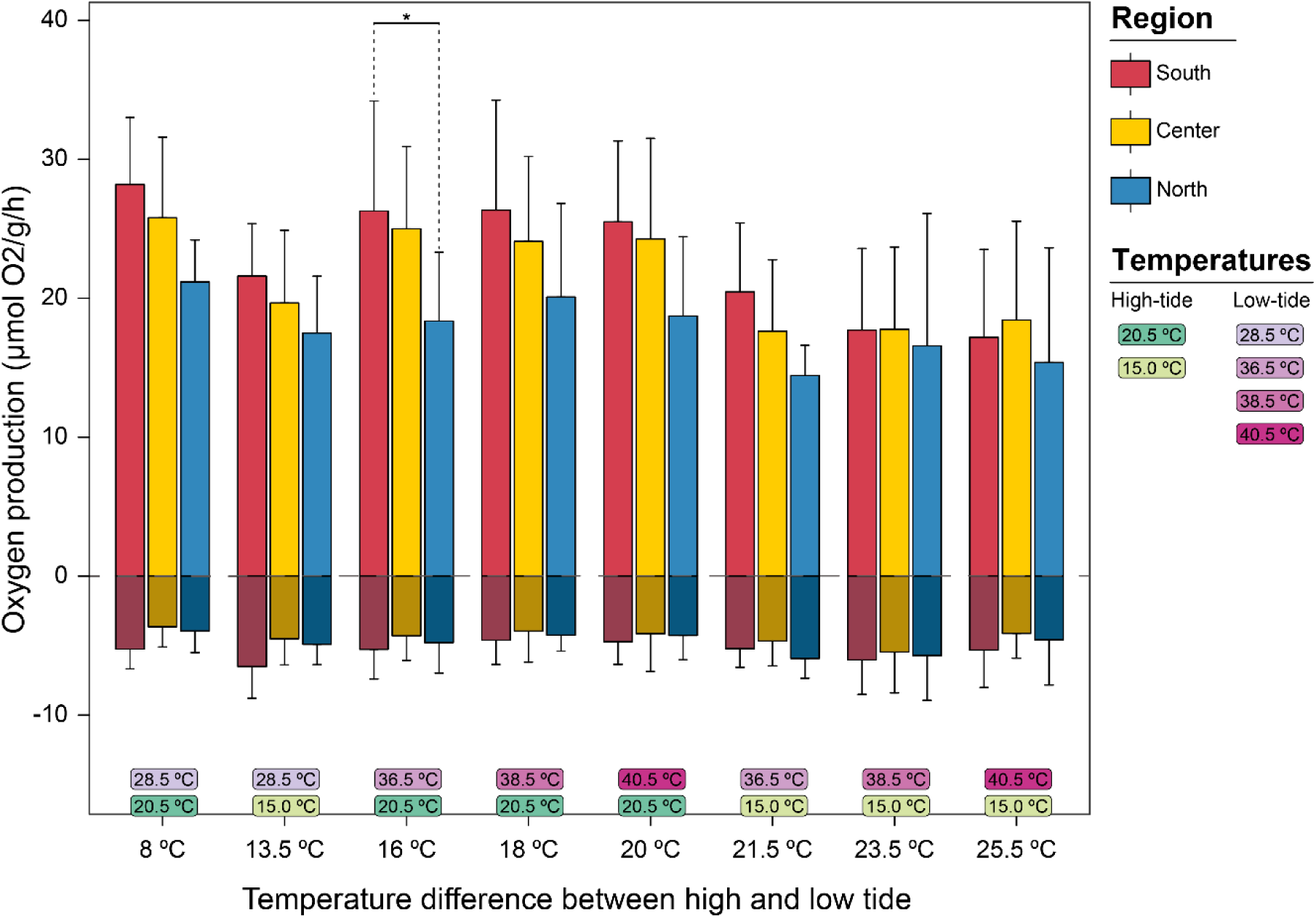
Maximum net primary production (Nppmax; µmol O2 g-1 h-1) and maximum respiration rate (Rmax; µmol O2 g-1 h-1) (mean + SD; n = 24 per region for South and Center; n=12 for North) after the recovery period (on the day 20). Regions are color-coded as blue (north populations), yellow (center populations), and red (south populations). Nppmax is shown as positive values, while Rmax is shown as negative values (and indicates consumption). The treatments are arranged from left to right based on the absolute temperature difference between lower temperatures at high tide and peak heating temperatures at low tide. Treatment temperatures are represented in shades of green (high-tide temperatures) and pink (low-tide temperatures). Statistical analysis focused only on pairwise comparisons between regions within the same treatment. Significance levels: * p < .05, ** p < .01, *** p < .001.

